# A well-controlled BioID design for endogenous bait proteins

**DOI:** 10.1101/427807

**Authors:** Giel Vandemoortele, Delphine De Sutter, Aline Moliere, Jarne Pauwels, Kris Gevaert, Sven Eyckerman

## Abstract

The CRISPR/Cas9 revolution is profoundly changing the way life sciences technologies are used. Many assays now rely on engineered clonal cell lines to eliminate overexpression of bait proteins. Control cell lines are typically non-engineered cells or engineered clones implying a considerable risk for artefacts because of clonal variation. Genome engineering can also transform BioID, a proximity labelling method that relies on fusing a bait protein to a promiscuous biotin ligase, BirA*, resulting in the tagging of vicinal proteins. We here propose an innovative design to enable BioID for endogenous proteins wherein we introduce a T2A-BirA* module at the C-terminus of endogenous p53 by genome engineering, leading to bi-cistronic expression of both p53 and BirA* under control of the endogenous promoter. By targeting a Cas9-cytidine deaminase base editor to the T2A auto-cleavage site, we can efficiently derive an isogenic population expressing a functional p53-BirA* fusion protein. Using quantitative proteomics we show significant benefits over classical ectopic expression of p53-BirA*, and we provide a first well-controlled view on the proximal proteins of endogenous p53 in colon carcinoma cells. This novel application for base editors expands the CRISPR/Cas9 toolbox and can be a valuable addition for synthetic biology.

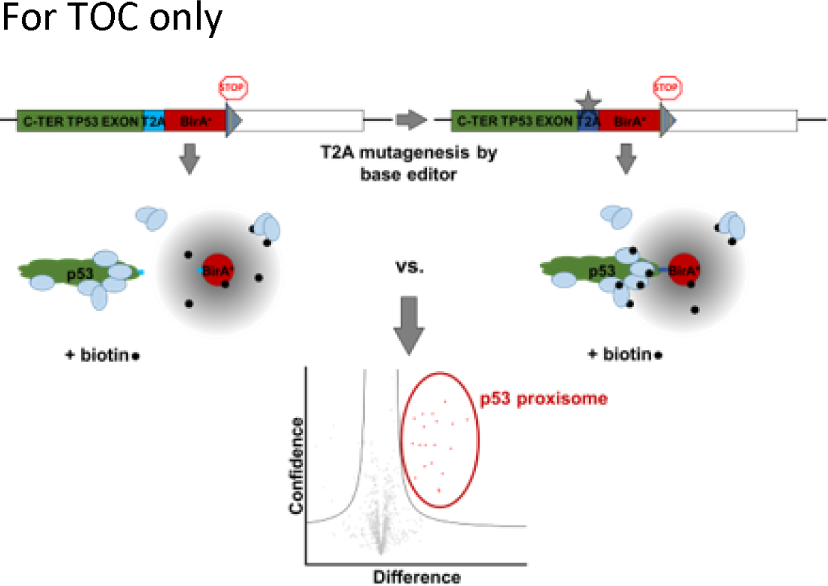

## INTRODUCTION

Elucidating the composition and function of protein complexes is crucial for advancing our understanding of key cellular processes, both in health as well as disease ^1^. Fuelled by an ever-growing arsenal of so-called interactomics technologies ^2^, vast amounts of data are now available for further exploration ^3, 4^. With expanding lists of interactors for every protein challenged, there is a growing consensus in the field that many reported interactions are highly contextual or spatially restricted ^5^. In addition, it became apparent that approaches in which a bait protein is overexpressed lack the subtlety that is needed to further dissect protein-protein interactions and assess physiological functions of individual interactions ^6^. Indeed, protein overexpression can have a detrimental effect on protein localization and folding, and distort downstream signalling ^5, 7^. Protein overexpression also tends to mask complex feedback control mechanisms affecting a protein at physiological levels and generally increases the overall stress status of a cell ^8, 9^. Several efforts have been made to avoid bait protein overexpression, with the use of primary antibodies to pull-down endogenous protein complexes as prime example ^10, 11^. In reality, the specificity and the sensitivity of antibodies are often inadequate for studying endogenous protein complexes in a reliable manner and can therefore lead to problems with reproducibility ^12^. Despite improved methodologies it remains challenging to generate specific antibodies with favourable affinities for each protein ^13, 14^ especially when one wants to specifically study mutant proteins bearing amino acid substitutions or truncated proteins originating from, for instance, alternative translation start sites (i.e. proteoforms ^15^). For such situations, Flp-In™ cell lines allowing tunable expression provide a valuable alternative ^16^. However, this requires thoughtful and precise fine-tuning of expression levels to mimic as closely as possible physiological protein levels ^17^. Artificial chromosomes have also been used to express transgenes at physiological levels ^18^.

A technology that is gaining popularity to map protein interactomes is BioID. This method relies on a promiscuous biotin ligase (BirA*) fused to a bait protein of interest ^19^. BirA* is a mutagenized *E. coli* biotin ligase (carrying the R118G mutation) that no longer displays substrate specificity. Upon expressing a bait-BirA* fusion protein, proximity-dependent biotinylation occurs of proteins that are near-neighbours of this bait protein in the cell. As endogenous biotinylation is a rare and highly specific modification in mammalian cells ^20, 21^, biotinylated prey proteins are easily discriminated after purification by streptavidin matrices and mass spectrometry (MS) analysis. Moreover, the need to keep protein complexes intact during lysis is removed since the labelling of prey proteins occurs in intact cells before the homogenization step. Since its initial publication in 2012, BioID has been used to study proximal and interacting proteins (i.e. the proxisome) of bait proteins in several organisms and cell types producing datasets complementary to affinity purification – MS (AP-MS) approaches ^22^. In bait overexpression experiments, as controls, BirA* is often expressed alone or fused to irrelevant baits (e.g. eDHFR, eGFP), although cellular localizations and expression levels might differ between BirA*-fusion proteins and BirA* alone, which therefore confound results ^19^.

Bi-cistronic expression systems can provide a way to ensure similar expression levels for a bait protein and a detached BirA* module. 2A peptides cause ribosomal skipping during translation and recommencement of translation immediately afterwards ^23^. 2A peptides display increased efficiency when compared to Internal Ribosome Entry Site (IRES) sequences and are only 18 amino acids in length, which is an additional advantage when considering genome engineering approaches ^24^.

Since the emergence of CRISPR/Cas9, many applications have been published with several of them outgrowing classical genome engineering ^25^. A recent application is on base editing in which Cas9 is fused to a cytidine deaminase. This enzyme converts cytosine into uracil which has the base-pairing properties of thymine ^26^ and thus introduces mutations. The base editing activity can be targeted to confined sequence windows of approximately five nucleotides by loading Cas9 with a specific guide RNA (gRNA). As base editing is typically performed by nickase Cas9 (nCas9) or catalytically inactive (dead, dCas9) fusion constructs, no DNA double strand breaks (DSBs) are introduced, implying that minimal numbers of insertion or deletion (indel) mutations are generated in this process. To avoid base excision repair which would revert the introduced mutation, extra moieties were added to these base editors, such as an uracil DNA glycosylase inhibitor (UGI) ^27^. Up to now, base editing applications mainly focussed on correcting disease mutations, on genetic diversification using tiled gRNA libraries ^27, 28, 29^, or on introducing premature stop codons as alternative to classic knock-out strategies ^30^.

p53 is arguably the most studied protein in health and disease ^31, 32^. Historically dubbed the guardian of the genome ^33^, the encoding gene is mutated in 50% of all reported human cancer cases ^34^, making it an attractive starting point for the development of anticancer strategies ^35^. Because of the long-lasting research focus on p53, the current interactome of p53 consists of over 1000 protein interaction partners (BioGRID, version 3.4 ^3^). p53 acts as a homotetrameric, sequence-specific transcription factor capable of inducing large gene networks comprising hundreds of target genes ^36, 37^. Known pathways regulated by p53, either by transactivation or protein interactions, include DNA damage repair, cell cycle arrest, senescence and apoptosis following exposure to a plethora of stress conditions such as DNA damage, telomere erosion, hypoxia, replication stress and oncogene activation. More recently, additional processes under p53 control such as autophagy, metabolism and ferroptosis were revealed illustrating the need for continued research on this protein ^38^.

In this report, we use a novel knock-in strategy to enable BioID experiments for endogenous p53 in HCT116 colon carcinoma cells as classic overexpression experiments in this cell model gave suboptimal results. In our knock in strategy, a control cell line was first generated by introduction of a T2A-BirA* cassette at the C-terminus of endogenous p53 using CRISPR/Cas9 genome engineering. These cells express the BirA* protein at equal levels to p53 and thus provide a good system to assess biotinylation background. Targeted inactivation of the T2A peptide sequence by a Cas9 cytidine deaminase fusion (BE3) results in the generation of a cell line that expresses a functional p53-BirA* fusion protein from the endogenous promoter allowing proximal protein mapping for endogenous p53. This reduces genome engineering efforts considerably and eliminates artefacts due to clonal variation when a selection step is added in the protocol. Furthermore, the concept of auto-cleavage switching in fusion proteins can find applications in synthetic biology.

## EXPERIMENTAL PROCEDURES

### Cell culture

HCT116 colon carcinoma cells were purchased from ATCC (ATCC CCL-247) and cultured in McCoy’s 5A modified medium (Gibco, 22330070). Genomically characterized HEK293T embryonic kidney cells ^39^ were cultured in DMEM (Gibco, 61965026) supplemented with fetal bovine serum (FBS) (Gibco, 10270106) to a final concentration of 10%, 12500 units penicillin-streptomycin (Gibco, 15070063) and HEPES (Gibco, 15630056) to a final concentration of 10 mM. Growth medium was switched to McCoy’s 5A modified medium or DMEM supplemented with 10% dialyzed FBS (Gibco, 26400044) for HCT116 2 weeks prior to seeding cells for a BioID experiment. Cells were passaged when cultures reached 70% confluence. All cell lines were deemed clear of mycoplasma by using a mycoplasma PCR detection kit (Minerva Biolabs, MIN-11-1100).

### Plasmids and cloning procedures

The BioID biotin ligase BirA* was transferred from the pcDNA3.1 mycBioID vector (Addgene #35700) by PCR to generate the pAav-TP53-T2A-BirA* targeting construct (available via Addgene, #115652). All primers used in this manuscript can be found in table S1. Homology regions were generated by PCR on genomic HCT116 DNA as described before ^40, 41^. The pMet7-FLAG-PQS1-TP53-T2A-BirA*-Myc construct was generated by standard restriction enzyme cloning procedures using the FLAG-PQS1-p53 expression vector published earlier ^40^. This construct was further mutagenized by PCR to obtain a construct bearing an inactivated T2A sequence for a BioID transient transfection experiment. A puromycin selection cassette was incorporated in the BE3 (Addgene, plasmid #73021) vector for endogenous mutagenesis purposes (BE3_puro_) using a linear dsDNA fragment (IDT DNA Technologies, gBlock). gRNA sequences incorporated in a U6-driven expression cassette were ordered as gBlocks (Table S2) prior to blunt-ended ligation in a pCR-Blunt vector (Thermo Fisher Scientific, K2750).

### CRISPR/Cas9 genome engineering

HCT116 *TP53*^+/T2A-BirA*^ cells were generated using a combination of CRISPR/Cas9 and recombinant Adeno-associated virus (rAAV)-mediated template delivery as described before ^40, 41^. Briefly, a *TP53* C-terminal targeted gRNA (5’ ACGCACACCUAUUGCAAGCA 3’) was cloned in a Cas9 expression construct (Addgene #48139) as described by Ran *et al*. ^42^. Cells were seeded in a 24-well plate 24 h before transfection with the Cas9 plasmid using FugeneHD (Promega, E2311). 48 h after transfection, cells were infected with targeting construct packaged in rAAV. Neomycin selection (Gibco, 11811031) was performed for two weeks starting 24 h after infection. Afterwards, cells were manually diluted and seeded as single cells in 96-wells for further clonal expansion until on average, clones reached 70% confluence at which point they were split into a lysis plate and culture plate. Screening of clonal populations in the lysis plate was based on junction PCR screening using GoTaq G2 Hot Start polymerase (Promega, M7405). Input lysates for PCR were obtained using DirectPCR lysis reagent (Viagenbiotech, 301C). Positive clones were seeded at 10^3^ cells per 96-well 24 h prior to treatment with 2.5 μM TAT-Cre (Excellgen, EG1001). 24 h after treatment, the medium was aspirated and replaced with fresh growth medium. Four days after TAT-Cre treatment, cells were subjected to a second round of single cell seeding and PCR screening. Clones showing aberrant growth rates or morphology were omitted for further characterization. PCR-positive clones were subjected to Western blot using 1/1000 diluted anti-p53 antibody (Calbiochem, OP43T) and near-infrared fluorescent secondary antibodies (LI-COR). Additionally, Southern blot analysis was performed to verify single genomic integration of the floxed neomycin resistance cassette. [α- ^32^P] Southern blot probe was generated as described before. ^41^ For every retained clone, engineered loci were verified using Sanger sequencing as described earlier. Primers used for screening, sequencing and generation of the Southern blot probe can be found in table S1.

### T2A mutagenesis

7.2×10^5^ HEK293T cells were seeded in 6-well plates the day prior to transfection. A total amount of 800 ng DNA was mixed with 4 μl polyethylenime (PEI) for every transfection condition and incubated for 10 min at room temperature. Meanwhile cells were placed on 1.5 ml DMEM medium supplemented with 2% FCS. 6 h after addition of the transfection mix, the medium was removed and replaced by DMEM with additives after washing once with Phosphate Buffered Saline (PBS) (Gibco, 14190094). 72 h post transfection cells were washed with PBS before detachment using a cell scraper (Greiner, 541070) and lysis in 2X SDS-PAGE XT sample buffer (Bio-Rad, 1610791) and XT reducing agent (Bio-Rad, 1610792) by heating for 10 min at 95°C. Afterwards, 100 μl MilliQ water was added to dilute samples prior loading on a QIAshredder homogenizer spin-column (Qiagen, 79656) and running them on a Criterion XT 4-12% BisTris gel (Bio-Rad, 3450123). After blotting, the PVDF membrane was incubated with anti-FLAG (Sigma, F1804) and anti-Myc (in-house produced) primary antibodies and detected with LI-COR near-infrared fluorescent secondary antibodies. For endogenous mutagenesis, BE3_puro_ vector and gRNA plasmids were transfected using FugeneHD in HCT116 *TP53*^+/T2A-BirA*^ cells. 24 h after transfection, 2 μg/ml puromycin was added to the growth medium for 48 h to enrich for BE3_puro_ transfected cells prior to expansion for proteomics experiments.

### Illumina targeted sequencing

For targeted sequencing, subconfluent 25 cm^2^ bottles containing HCT116 *TP53*^+/T2A-BirA*^ cells after BE3_puro_ treatment and subsequent enrichment were pelleted by centrifugation at 500 g for 5 min and washed once with PBS. To each pellet, 500 μl lysis buffer [20 mM Tris-HCI pH 7.5, 5 mM EDTA, 150 mM NaCl, 400 U Proteinase K (Sigma, P4850), 0.2% SDS] was added and incubated for 16 h at 37°C. Afterwards, genomic DNA was obtained by phenol-chloroform extraction and ethanol precipitation. Next, the T2A locus was amplified and provided with Nextera overhangs by PCR using Herculase polymerase (Agilent, 600675). Sequencing was performed using the MiSeq Nano v2 kit (Illumina, MS1022002) by the VIB Nucleomics core facility (http://www.nucleomics.be/). FASTQ files were uploaded to CRISPRESSO ^43^ to analyse results. The CRISPRESSO allele frequency output file (Table S3) was used as input to generate the sequence logo using the online tool WebLogo ^44^.

### Nuclear and cytoplasmic extraction

For every cell line, cells were cultured in McCoys medium supplemented with a final concentration of 10% dialyzed FBS as described above. 1 mM doxorubicin (Sigma, D1515) and 50 μM biotin (Sigma, B4639) was added to the growth medium 24 h prior to the start of the experiment at which 7×10^6^ cells were pelleted by centrifugation at 500 g for 5 min. After washing once with ice-cold PBS and centrifugation at 500 g for 5 min, the supernatant was aspirated from the cell pellet. Extraction was done using the NE-PER kit (Thermo Fisher Scientific, 78833) according to the instructions provided by the manufacturer. In brief, 500 μl ice-cold CER 1 buffer was added to every cell pellet before resuspension by vortexing. After 10 min incubation on ice, 27.5 μl ice-cold CER 2 buffer was added to the mix and samples were centrifuged at 16000 g on 4°C for 5 min. Supernatant resulting from this centrifugation step was stored as cytoplasmic extract. The pellet fraction was resolved in 250 μl ice-cold NER and vortexed vigorously every 10 min for 15 s during an incubation period on ice for a total of 40 min. Afterwards, nuclear extract was obtained by centrifugation at 16000 g for 10 min at 4°C and the supernatant containing this nuclear extract was transferred to a new tube.

In parallel, 10^6^ cells were washed once with PBS prior to resuspension in 150 μl ice-cold RIPA lysis buffer [50 mM Tris-HCI pH 7.5, 150 mM NaCl, 1% NP-40, 1 mM EDTA, 1 mM EGTA, 0.1% SDS, 0.5% sodium deoxycholate (DOC) and complete protease inhibitor cocktail (Roche, 11873580001)]. Following 10 min incubation on ice, lysates were centrifuged at 16000 g for 15 min at 4°C and the supernatant was transferred to a new tube (= total cell lysate sample). For every fraction, the protein concentrations as determined by a Bradford assay and 40 μg was loaded on ExpressPlus SDS-PAGE (GenScript, M42015) and afterwards blotted on a PVDF membrane. α-tubulin was used as cytoplasmic marker (Sigma, T5168).

### BioID affinity purification

For every sample three 145 cm^2^ culture dishes (Nunc) were seeded with 4.5×10^6^ HCT116 *TP53*^+/T2A-BirA*^ cells cultured on McCoy’s 5A modified medium supplemented with dialyzed FBS. 24 h after seeding 1 μM doxorubicin was added. To every culture dish biotin was added to a final concentration of 50 μM at the start of the treatment. 24 h after treatment the growth medium was replenished to deprive free biotin from the culture. 3 h later, the growth medium was aspirated and cells were washed once with cold PBS before detachment in PBS using a cell scraper. After pelleting cells by centrifugation for 5 min at 500 g on 4°C, a second PBS washing step occurred prior to cell lysis in a 15 ml tube. 2 ml of ice-cold RIPA lysis buffer containing 250 U of benzonase (Sigma, E1014) was added to each pellet. Lysis was obtained by incubation with agitation for 1 h on 4°C prior to sonication (30% amplitude, 5 × 6 s burst, 2 s interruption) on ice. The insoluble fraction of the lysate was removed by centrifugation for 15 min at 16000 g on 4°C and the supernatant was transferred to a new 15 ml tube. The total protein concentration was determined by the Bradford protein assay to normalize input material to the maximum shared total protein amount throughout the samples. For every sample, 90 μl Streptavidin Sepharose High Performance bead suspension (GE Healthcare, 17-5113-01) was used for enrichment of biotinylated proteins. Beads were pelleted by centrifugation at 500 g for 1 min and washed once in wash buffer (50 mM Tris-HCI pH 7.5, 150 mM NaCl, 1% NP-40, 1 mM EDTA, 1 mM EGTA, 0,1% SDS) prior to addition to the lysate. Samples were incubated for affinity purification on a rotator at 4°C for 3 h. Afterwards, beads were recovered by centrifugation of the samples at 500 g for 1 min at 4 °C and aspirating the supernatant. Beads were washed three times in wash buffer, twice in 50 mM ammonium bicarbonate pH 8.0 and once with 20 mM Tris-HCI pH 8.0, 2 mM CaCI_2_ prior to resuspension in 20 μl 20 mM Tris-HCI pH 8.0. Trypsin digest occurred overnight by addition of 1 μg trypsin (Promega, V5111) to each sample. Digestion mixtures were depleted of beads upon centrifugation at 500 g for 1 min, after which the supernatant was transferred to a mass spectrometry vial. To ensure complete digestion, peptide mixtures were incubated with an additional 500 ng trypsin for 3 h before addition of formic acid to a final concentration of 2%. All experiments were performed in biological triplicate for downstream label-free quantitative proteome analysis.

For transient transfection BioID experiments, 8.5×10^6^ HCT116 cells were seeded in a 56 cm^2^ culture dish (Nunc) and transfected the day after with pMet7-FLAG-PQS1-TP53-T2A-BirA*-Myc, the pMet7-FLAG-PQS1-TP53-MUTT2A-BirA*-Myc or pSV-SPORT mock vector using FugeneHD. 24 h post transfection, cells were transferred to 3 T145 cm^2^ dishes per sample. 24 h after transfer, 1 μM doxorubicin and 50 μM biotin were added 24 h prior to harvest. Affinity purification of biotinylated proteins was performed as described above.

### LC-MS/MS instrument analysis

Peptide mixtures were analyzed by LC-MS/MS on an Ultimate 3000 RSLC nano LC (Thermo Fisher Scientific) in-line connected to a Q-Exactive mass spectrometer (Thermo Fisher Scientific). The peptides were first loaded on a trapping column (made in-house, 100 μm internal diameter (I.D.) × 20 mm, 5 μm beads C18 Reprosil-HD, Dr. Maisch, Ammerbuch-Entringen, Germany). After flushing the trapping column, peptides were loaded in solvent A (0.1% formic acid) on a reverse-phase column (made in-house, 75 μm I.D. × 250 mm, 3 μm Reprosil-Pur-basic-C18-HD beads packed in the needle, Dr. Maisch, Ammerbuch-Entringen, Germany) and eluted by an increasing concentration solvent B (0.1% formic acid in acetonitrile) using a linear gradient from 2% solvent B up to 55% solvent B in 120 min, followed by a washing step with 99% solvent B, all at a constant flow rate of 300 nl/min. The mass spectrometer was operated in data-dependent acquisition (DDA), positive ionization mode, automatically switching between MS and MS/MS acquisition for the 5 most abundant peaks in a given MS spectrum. Source voltage was set at 3.4 kV, with a capillary temperature of 275°C. One MS1 scan (m/z 400-2000, AGC target 3×10^6^ ions, maximum ion injection time 80 ms), acquired at a resolution of 70000 (at 200 m/z), was followed by up to 5 tandem MS scans (resolution 17500 at 200 m/z) of the most intense ions fulfilling predefined selection criteria (AGC target 5 × 10^4^ ions, maximum ion injection time 80 ms, isolation window 2 Da, fixed first mass 140 m/z, spectrum data type: centroid, underfill ratio 2%, intensity threshold 1.3×E^4^, exclusion of unassigned, 1, 5-8 and >8 positively charged precursors, peptide match preferred, exclude isotopes on, dynamic exclusion time 12 s). HCD collision energy was set to 25% normalized collision energy and the polydimethylcyclosiloxane background ion at 445.120025 Da was used for internal calibration (lock mass).

### Mass spectrometry data processing and interpretation

Obtained Xcalibur Raw Files (.raw) were analyzed using MaxQuant and maxLFQ algorithms (MaxQuant version 1.5.8.3) ^45, 46^. Spectra were searched against the human UniProt sequence database (as available on 01/09/2017). Additional FASTA files for BirA (ID Z00001), BirA* (ID Z00002) and T2A (ID Z00003) sequences were included in the searches. Methionine oxidation, N-terminal acetylation and lysine biotinylation were set as variable modifications, with a maximum of 5 modifications per peptide. The minimum peptide length was set at 7 amino acids and a maximum peptide mass of 4600 Da was used. PSM, protein and site false discovery rates were set at 0.01. The minimum Label-Free Quantitation (LFQ) ratio count was 2 and the Fast LFQ option was disabled. 20 ppm and 4.5 ppm mass accuracies were used for the first and main search respectively. After completion of searches, LFQ intensities were loaded in Perseus (version 1.5.5.3) ^47^ for further analysis.

Samples were annotated on condition for analysis. Proteins only identified by site, reverse hits and Perseus contaminant proteins were removed from the data matrix. Retained LFQ intensities were transformed to log2 scale and three valid values in at least one of the two sample groups were needed in order for the protein to be retained for further analysis. Missing values were imputed from a normal distribution of intensities (0.3 width, 2.2 downshift). A two-sided t-test (0.05 FDR, 1000 randomizations) was performed to reveal differentially enriched proteins in the volcano plot. Optimal s0 values were calculated for all analyses using the SAM-test R package ^48^. Gene ontology (GO) analyses were performed using the database for annotation, visualization and integrated discovery (DAVID 6.8) ^49^. The mass spectrometry proteomics data have been deposited to the ProteomeXchange Consortium via the PRIDE ^50^ partner repository with the dataset identifier PXD011702.

## RESULTS

### BioID by forced expression of a p53-BirA* fusion protein in colon carcinoma cells

Proximity biotinylation is an exciting development of recent years to identify vicinal proteins of a target protein which includes the associated protein complexes ^19, 22^. While the downstream purification of biotinylated proteins of the *in vivo* biotinylation reaction is a powerful and sensitive procedure, great care should be taken with the design of the experiment and the levels of BirA* ^51^. Most BioID studies rely on inducible expression systems based on Flp recombinase integration (Flp-In™ T-REx™ system). This powerful system allows low expression which can be tuned to match endogenous levels. Despite being around for well over 10 years, only few engineered Flp-ln cell lines are available and virtually all BioID studies using this system are therefore performed in 293 cells ^51, 52, 53^

We set out to explore the so-called ‘proxisome’ of p53 in well-known and well-characterized colon carcinoma HCT116 cells. The model system has been used extensively for the study of p53 function, in part because of the availability of many knock out cell lines ^54^. We were faced with the lack of an inducible HCT116 cell line and we therefore decided to first explore an experiment with forced expression of a p53-BirA* construct. We transiently transfected HCT116 cells with a fusion construct that contains a C-terminal BirA* module cloned in frame through an inactivated T2A peptide that served as a long linker (p53-MUTT2A-BirA*). As control experiments we used either mock-transfected cells or cells transfected with a bi-cistronic p53-T2A-BirA* construct (Figure 1A and 1B). This last construct ensured equal levels of the free BirA* module in the control cells, providing a distinct advantage over expression of a free unfused BirA* protein which often requires tuning to obtain equal expression levels. We performed triplicate experiments to allow LFQ analysis using MaxQuant and Perseus workflows. Optimal s0 values for all proteomics experiments described in this work were derived using the SAM-test R package to enable unbiased assessment of the data ^48^. Using the mock control, an extensive list of significant proteins for p53-BirA* was obtained by LFQ analysis (Figure 1C). Of these, 16.7% were known p53 interaction partners in BioGRID. Some well-known interaction partners can be discerned in the list, but despite the comprehensive list, some key interaction partners such as MDM2 are lacking. This extensive list can be expected because the control conditions lack any BirA*. When assessing the T2A-BirA* control condition we found some distinct differences, pointing to removal of contaminants by the presence of BirA* in this control. However, the retrieved lists were still surprisingly long with 521 significant proteins (Figure 1D), of which 19.2% were known BioGRID p53 interaction partners. This large number of hits is likely due to the excessive levels of BirA*, resulting in much background biotinylation in both transient expression approaches. Levels of the fusion protein are well beyond the endogenous levels in the cells (Figure 1B). In addition, GO analysis on ‘molecular function’ showed very few links to p53 function, with ‘p53 binding’ only retrieved at position 59 and 82 for the mock and T2A-BirA* control respectively. GO analysis for ‘cellular component’ showed strong enrichment of cytoplasmic and membrane proteins (Table S4 and S5) which contradicts the predominant nuclear function of p53. Thus the results of these analyses hint to the presence of substantial amounts of biologically irrelevant, false positive prey proteins.

**Figure 1.**
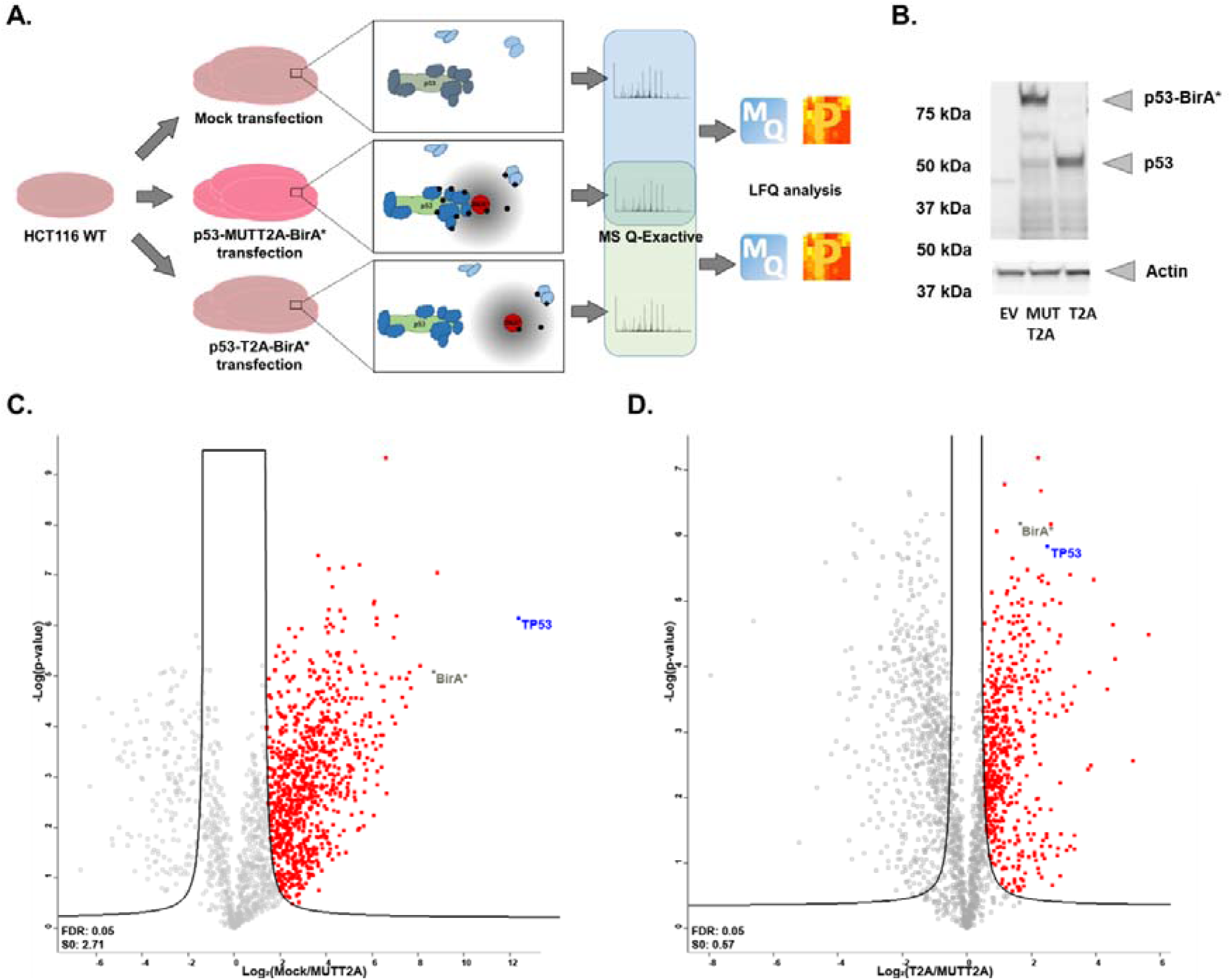
Transfection p53 BioID experiments. (A) Workflow used for transient expression LFQ proteomics experiments. All experiments were conducted in presence of 1 μM doxorubicin. Top panel: Control transfection with an empty vector. Middle panel: p53-MUTT2ABirA* transfection leading to biotinylation of vicinal p53 proteins. Bottom panel: p53-T2A-BirA* control transfection leading to biotinylation of background proteins. (B) Western blot for p53 in the transient expression experiment. Different lanes depict p53 protein levels in the empty vector transfection (EV), in the fusion p53-MUTT2ABirA* transfection (MUTT2A), and in the transfection with the bi-cistronic p53-T2A-BirA* (T2A) control. Loading control: anti-actin. (C) and (D) Volcano plots showing differential LFQ intensity levels (X-axis) and p-values (Y-axis) in transient transfection BioID experiments using a mock and BirA* control transfection respectively.

### Generation of a control cell line for BioID on endogenous p53

The long lists in the overexpression experiment are likely caused by the high expression levels of the bait protein. While this can be solved by introducing an inducible expression cassette containing a p53-BirA* fusion construct through lentiviral transduction ^55^, we decided to explore the exciting developments in the genome editing field to engineer the BirA* module in one allele of the *TP53* gene in HCT116 cells. We started by first creating a control T2A-BirA* cell line as we reasoned that the best way to ensure similar levels in the control condition is to use a bi-cistronic approach (Figure 2A). Similar BirA* control expression levels are also maintained when p53 protein levels are changed due to stress conditions on the cells (e.g. DNA damage). We used CRISPR/Cas9 to introduce a double stranded DNA (dsDNA) break in the last coding exon of *TP53* and provided a single strand DNA (ssDNA) repair template containing a T2A-BirA* module and a floxed expression cassette flanked by two homology regions. Generation of this control cell line was performed using a two-step engineering approach as described earlier (^40^, Figure S1). Clones bearing insertions were identified by PCR (Figure S2). In the second step, the floxed selection cassette is removed by CRE recombinase, again followed by PCR screening of single cell clones (Figure S3). Single integration for these clones was shown by Southern blot (Figure S4A), and by Sanger sequencing of both alleles ensuring an intact second allele (Supplementary Figure S4B and S4C). Because of current limits in knock-in efficiencies, single cell clones were selected and evaluated for the introduction of the knock-in construct and for CRE-based removal of the selection cassette.

**Figure 2.**
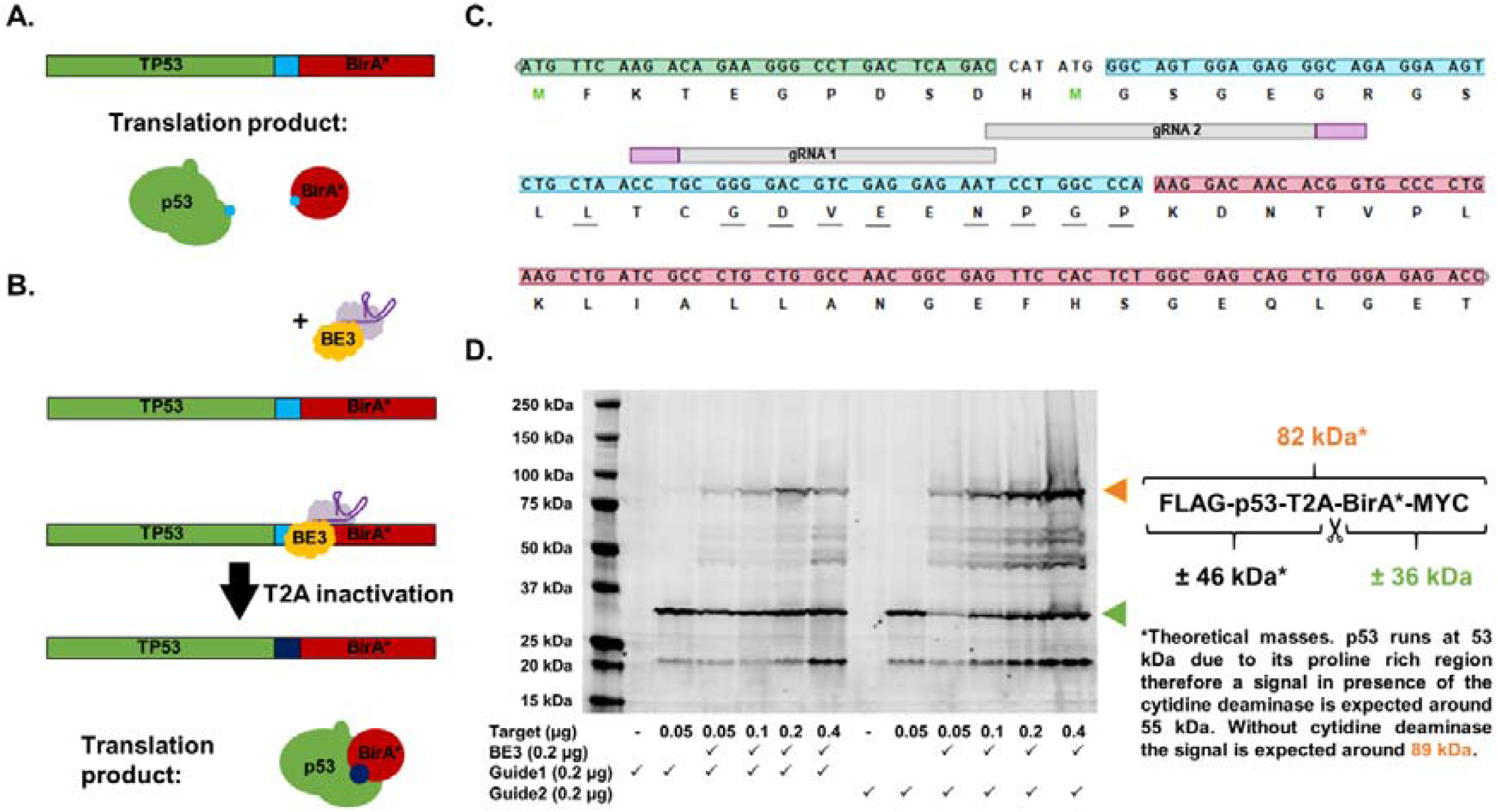
Targeted T2A inactivation. (A) Incorporation of a sequence coding for a T2A peptide (cyan) between the bait (p53) and BirA* will lead to translation of two separate proteins. (B) After transfection, the BE3 base editor will be guided to the T2A sequence by (a) specific gRNA(s) and will convert cytidines mainly to thymidines. Dark blue: inactivated T2A (MUTT2A) (C) Layout of the T2A region in the TP53-T2A-BirA* test construct. Conserved T2A residues are underlined. Green: p53 C-terminus; cyan: linker and T2A sequence; red: BirA* N-terminus; grey: gRNA used; magenta: PAM-motif. (D) Overexpression pilot experiment to assess T2A mutagenesis of a target plasmid. Cells were lysed 72 h after transfection with a MYC-tagged p53-T2A-BirA* construct, and loaded on SDS-PAGE for Western blotting using an anti-MYC antibody. Upon mutagenesis a band corresponding to the 82 kDa p53-MUTT2A-BirA* protein is visible. The two gRNAs targeting different residues of the T2A sequence were tested in parallel for potency (left: gRNA 1; right: gRNA 2). In both cases, expression of p53-BirA* could be observed at varying amounts of target concentrations. gRNA 2 was selected for further endogenous use.

### Deriving an experimental cell line using a base editor approach

An experimental cell line containing an engineered *TP53* locus with a C-terminal BirA* module can be obtained using a similar genome engineering approach as for the control cell line, but there are a number of things to consider. Firstly, the seeding, culturing and screening of single cell clones related to current knock-in efficiencies requires a significant effort effectively doubling genome engineering hands-on time to perform a BioID experiment. Secondly, the generation of the experimental cell line would also require two selection steps of single cell clones. This could introduce clonal variation and may introduce artefacts in the data. To solve these issues, we sought to derive an isogenic population from the control cell line wherein the T2A site would not be present. This can be obtained by another CRISPR/Cas9 knock-in approach using a template with an alternative linker (e.g. an inactive T2A site). However, the efficiencies currently obtained for knock-in are too low to allow such an approach and would still require clonal selection (see higher). As an alternate way to achieve T2A inactivation, we turned to a recently reported Cas9-cytidine deaminase base editor BE3 ^27^. Targeted inactivation of the 2A sequence would ensure restoration of bait-BirA* transgene expression (Figure 2B). 2A peptides have conserved amino acid residues that are critical for peptidyl transferase inhibition that causes ribosomal skipping (Figure 2C) ^56, 57^. Therefore, the targeted mutagenesis of one of these key residues should enable efficient abrogation of 2A ribosomal skipping activity. To explore this, we generated an expression vector that expresses a fusion construct of the tumor suppressor protein p53 linked to BirA* by a T2A sequence. Appropriate codon usage of the T2A sequence was verified to ensure the presence of several target nucleotides for BE3 (Figure 2C). Varying amounts of a FLAG-TP53-T2A-BirA*-MYC expression construct were co-transfected with the BE3 construct and T2A-specific gRNA. Two different gRNAs were assessed for base editing activity (Table S2). Conversion of the auto-cleavage site was clearly detectable by the presence of a high molecular weight band corresponding to a p53-BioID fusion construct on Western blot analysis (Figure 2D).

### T2A conversion after integration by genome engineering

A puromycin selection cassette was first cloned into the original BE3 vector allowing for efficient enrichment of transfected cells. This ensures that non-transfected cells, and thus non-edited cells, are efficiently removed in a short puromycin selection step. These non-edited cells would generate additional noise to downstream LFQ proteomics analysis ^45^. The gRNA directly targets the conserved amino acid residues on the T2A sequence while leaving p53 and BirA* intact (inactivated T2A is hereafter indicated as MUTT2A). When assessing p53 translation products before or after BE3 transfection, a clear signal for p53-MUTT2A-BirA* was observed in puromycin-resistant cells (Figure 3A), a result which was obtained for multiple clones. Notably, an additional band with an apparent molecular weight corresponding to WT p53 was also present. This extra band can be explained by the heterozygous knock-in of T2A-BirA*. In HCT116 *TP53*^+/T2A-BirA*^ the modified p53 allele has 22 extra amino acids due to translation of the GSG linker and the T2A peptide. This band could still be observed in the enriched cell population post-transfection, hinting at a portion of cells that did not undergo T2A sequence conversion. This partial conversion can be attributed to the intrinsic activity of BE3 that is below 100%. Furthermore, it is likely that not all editing will result in T2A inactivation as BE3 mutagenesis activity is centered around positions 4 – 8 in the protospacer sequence ^27^, therefore mutations might be introduced that have no impact on T2A activity or only attenuate its potency. To quantify the mutagenesis rate we performed next-generation sequencing on the resulting cell population showing a conversion rate over 40% (Figure 3B). In agreement with earlier reports on base editing, CRISPRESSO analysis showed that the rate of indel formation in the resulting population was negligible and the mutagenesis window was restricted to the predicted nucleotides (Figure S5).

**Figure 3.**
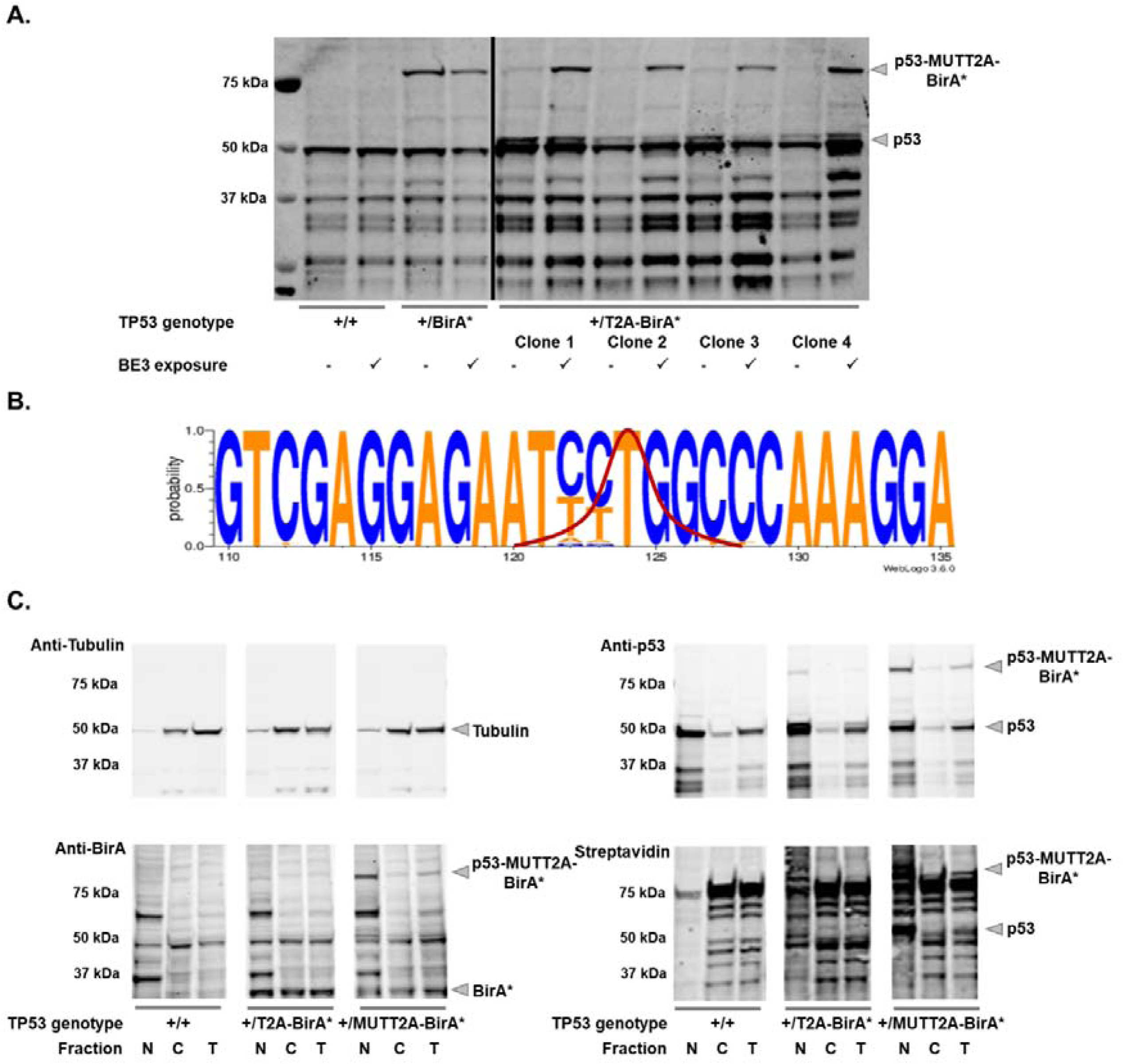
Endogenous T2A mutagenesis and cell line characterization. (A) Endogenous T2A mutagenesis. Cells were lysed 72 hours after BE3 transfection and loaded on SDS-PAGE for Western blot analysis using an anti-p53 antibody. p53 or p53-BioID expression is not influenced by BE3 expression in parental HCT116 cells or in HCT116 *TP53*^+/BirA*^ control cell lines. T2A inactivation could be observed for all four HCT116 *TP53*^+/T2A-BirA*^ clones tested resulting in expression of the p53-MUTT2A-BirA* fusion protein. Clone 4 was used for further experiments. (B) Sequence logo depicting mutation frequencies of BE3-treated cells. Paired-end sequencing was performed on the cell population after BE3 treatment and enrichment. The relevant T2A region is depicted in the sequence logo. C > T conversions could only be observed in the targeted residues. Red: expected BE3-induced mutation prevalence according to the position of gRNA 2. (C) Nuclear and cytoplasmic extraction experiments of doxorubicin-treated cell lines. p53 is mostly present in the nuclear extract of all cell lines. Both before and after BE3 treatment, BirA* shows an identical localization in the nucleus, in accordance with p53. In both populations, biotinylation is elevated when compared to parental HCT116 cells, with the majority of biotinylated substrates detected in the nuclear extract. N: nuclear extract; C: cytoplasmic extract; T: Total cell lysate.

Next to efficient T2A inactivation, proper localization of p53-MUTT2A-BirA* and BirA* is clearly crucial for the effectiveness and reliability of the system. Ideally, both the bait and the biotinylation activity of the engineered cell lines should follow the same subcellular distribution before and after BE3 treatment to provide relevant background protein signals. To evaluate this, we derived nuclear and cytoplasmic fractions of the engineered cells upon p53 activation by doxorubicin. We could observe a clear nuclear enrichment for p53 in both parental and engineered cells, indicative that proper p53 localization is not impaired by knock-in of T2A-BirA* or base editor treatment (Figure 3C). As expected, only in the BE3 treated engineered cells a strong signal for p53-T2A-BirA* could be observed in the nuclear fraction. Furthermore, assessment of the biotinylation pattern showed an intense laddering pattern in engineered cells when compared to the parental HCT116 cells (Figure 3C). Intriguingly, no biotinylated free BirA* could be observed when probing with streptavidin. Both before and after BE3 treatment, biotinylation was mainly detectable in the nuclear fraction, which indicates that the engineered cells with cleaved BirA* (p53-T2A-BirA*) provide a good control condition. Additionally, biotinylated WT p53 can be clearly observed in BE3 treated cells, hinting that fusion to BirA* is not affecting the oligomerization properties of p53 (Figure 3C).

### Conversion of inserted T2A-BirA* as a potent tool to study the p53 proxisome

We then performed endogenous BioID experiments using the engineered and converted HCT116 *TP53*^+/MUTT2A-BirA*^ cells. Experiments were done in triplicate to allow MaxQuant LFQ analysis ^45^. Cells were harvested and lysed after provoking a p53 response by doxorubicin. Puromycin-selected HCT116 *TP53*^+/2A-BirA*^ cells transfected with BE3 in absence of T2A-directed gRNA served as control condition (Figure 4A). Following a two-sided t-test (FDR: 0.05) we observed 207 potential p53 interactors for the mutagenized cells (Figure 4B), resulting in a high-quality proxisome dataset for p53. String pathway analysis showed a highly interconnected network around p53 with complexes involved in transcription and chromatin remodelling (Figure S6). Among the most differential proteins is p53 (p-value <0.00001), supporting that T2A is mutagenized in a way that abrogates ribosomal skipping by the peptide. In the control condition treated with doxorubicin, p53 was detected at lower levels, hinting at rare events whereby the T2A peptide did not cause interruption of translation in accordance with the Western blot analysis (Figure 3A). Next to p53, 75 (36.2%) known direct p53 interaction partners were identified including several hallmark interactors such as MDM2 ^58^, TP53BP1 ^59^, EP400 ^60^, and Sin3B ^61^. Noteworthy, we also detected proteins that affect the post-translational (PTM) signature of p53 such as the phosphatase PPP1R10 ^62^. Several proteins that could not directly be linked to p53 have been already implied in DNA damage pathways such as DMAP1 ^63^ and EPC1 ^64^. For virtually all significantly enriched proteins, multiple unique peptides were detected, enabling a robust protein inference (Table S6). GO analysis showed highly specific enrichment for p53-related ‘cellular component’ and ‘molecular function’ terms (Table S7). ‘p53 binding’ can be found at position 12 with a p-value of 3.93×10^−10^. These results support the concept that BioID on the endogenous protein level can be exploited to further characterize dynamic p53 interactions in a physiologically relevant context. Interestingly, The BirA* protein was enriched in the MUTT2A condition (Figure 4B). This is remarkable, as we observed that the expression level of free BirA* was comparable or even decreased before and after BE3 treatment in the HCT116 *TP53*^+/T2A-BirA*^ (Anti-BirA Western blot, Figure 3C). This observation can probably be attributed to the lack of biotinylation of free BirA* in both engineered cell lines (Western blot with streptavidin, Figure 3C). Thus, BirA* is only purified as a part of the p53-MUTT2A-BirA* fusion protein resulting in the observed difference upon MS analysis.

**Figure 4.**
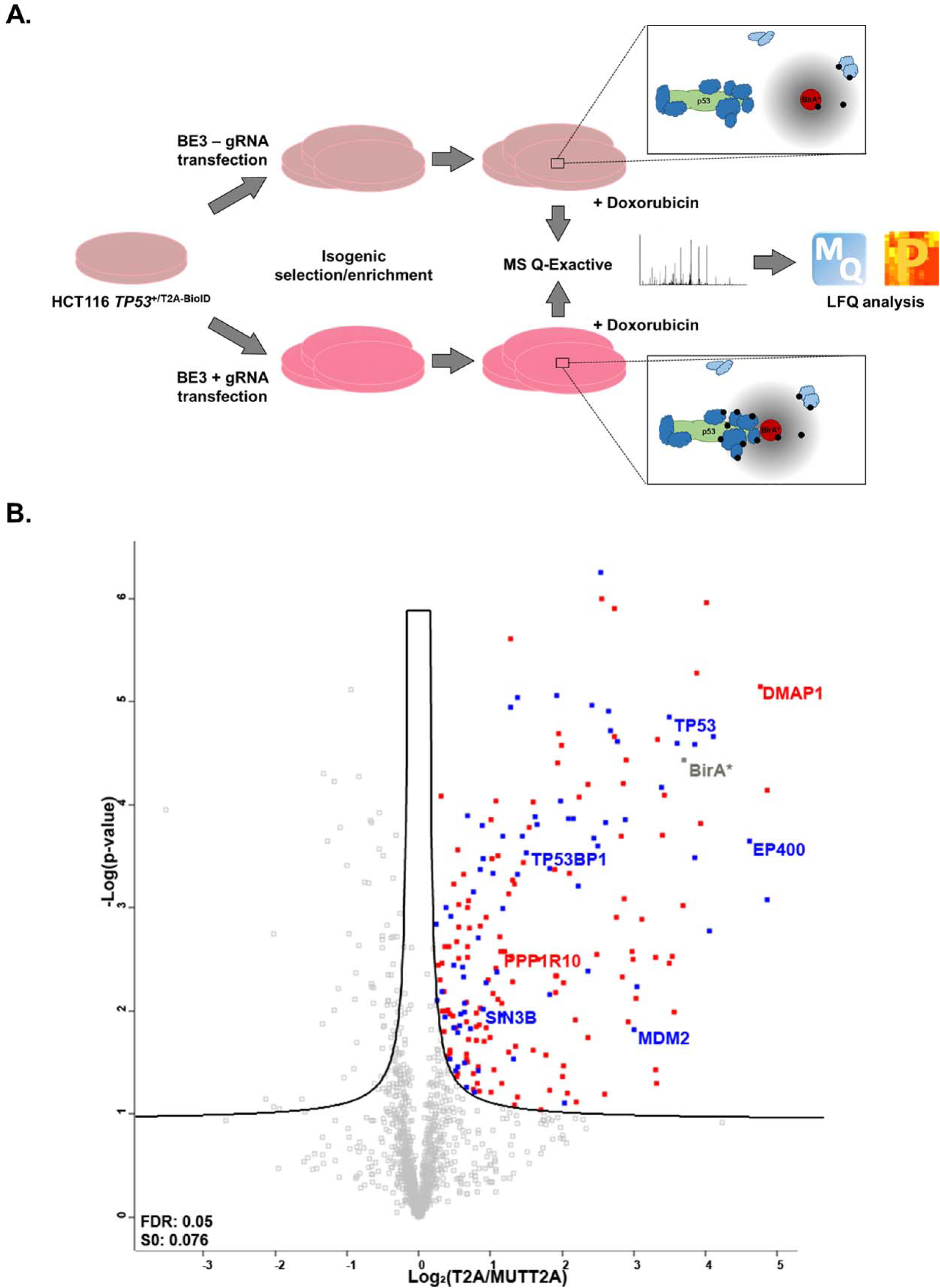
Mapping the endogenous p53 proxisome using BioID. (A) Proteomics workflow used for HCT116 *TP53*^+/T2A-BirA*^. Experiments were done in triplicate to support LFQ analysis. The puromycin resistance cassette in the BE3_Puro_ vector was used for enrichment of the transfected cells. Cells were allowed to recover prior to doxorubicin treatment to activate p53. Zoom: graphic representation of the promiscuous biotinylation in HCT116 *TP53*^+/T2A-BirA*^ (top) and HCT116 *TP53*^+/MUTT2A-BirA*^ (bottom). Blue: p53-proximal proteins. Black: biotin. (B) Volcano plot showing the differential LFQ intensity levels (X-axis) and the p-values (Y-axis) for HCT116 *TP53*^+/T2A-BirA*^ cells before and after T2A conversion in presence of doxorubicin. Known p53 interactors indexed in BioGRID are highlighted in blue. Significant hits discussed in the main text are labeled.

To show the benefit of using a converted T2A design strategy, we also compared non-engineered WT control cells (transfected with BE3 and selected on puromycin) to the p53-MUTT2A-BirA* cells as this would mimic a simpler design strategy (Figure S7A). This analysis revealed 418 differential proteins with a clear enrichment of RNA-binding proteins and chaperones (Figure S7B; Table S8) with ‘p53 binding’ found on position 18 with a p-value of 6.11×10^−8^. When compared to the T2A-BirA* control condition, 233 proteins were exclusively significant for the analysis using transfected HCT116 WT cells (Figure S7C). A part of this extended protein list is also found in the BioGRID database as interactors of p53. Strikingly, when these additional p53 BioGRID hits were assessed for ‘cellular component’ terms they were mainly assigned as cytoplasmic proteins (188 proteins; p-value 7.43×10^−14^), and their ‘molecular function’ was linked to functions such as RNA-binding (57 proteins; p-value 2.61×10^−20^), ribosomal constituents (31 proteins; p-value 1.89×10^−14^) and poly(A) RNA binding (148 proteins; p-value 6.17×10^−70^; Table S8). While these RNA-binding proteins and chaperones can be *bona fide* interaction partners for p53, they likely turn up with many different bait proteins (or directly with BirA*). Consequently, proteins with functions related to RNA are typically enriched in contaminant repositories for affinity purification – mass spectrometry experiments ^65^.

## DISCUSSION

Since its initial emergence, BioID has been widely adopted by the proteomics community to detect protein-protein interactions and vicinal proteins because of its distinct advantages over classic AP-MS ^22^. However, virtually all efforts rely on ectopic (over)expression of bait proteins fused to BioID ^22^. To our knowledge, there is currently only one report where BioID is performed at the endogenous level ^66^. The endogenous application of BioID solves two important issues in typical MS-based interactomics approaches. Firstly, the need for lysis is obviated since prey labelling occurs in intact cells before the homogenization step, and secondly, overexpression artefacts are removed by using the endogenous bait protein. This last issue is supported by our experiments with ectopic expression of p53-BirA* wherein we clearly show extensive lists of candidate proximal proteins pointing to the presence of many contaminants. Coupling expression levels of both bait-BirA* and control BirA* might be of particular relevance for proteins, such as p53, that have varying expression levels depending on the cellular state. One longstanding question in the p53 field is how p53 is able to elicit different responses to separate stimuli or stress conditions in a cell. We believe that future studies with BioID at the endogenous level may reveal some of the subtle interaction changes or altered feedback loops that are potentially overlooked in overexpression studies, thereby providing missing pieces to the p53 puzzle. Furthermore, the majority of p53 research has been based on overexpression experiments and therefore PPI databases may be riddled with interactions that are spatially and physiologically not possible in an intact cell. Although highly specific anti-p53 antibodies are commercially available for conducting endogenous immunoprecipitation MS experiments, these will always have the drawback that protein complexes need to be kept intact during purification. While crosslinking of protein complexes may improve conservation during purification, crosslinking is prone to inter-experimental variation in efficiency, suffers from ambiguous experimental data and can lead to problematic identification of crosslinked peptides by search algorithms ^67^.

We engineered a single cell line that can be used both as a control and an experimental cell line without the need to go through an additional intermediary single cell stage. Our approach relies on mutagenesis of a T2A sequence incorporated as a linker between a bait protein of interest and a functional tag sequence, here BirA*. By mutagenizing the T2A site, ribosomal skipping is prevented and a functional fusion construct is generated. We showed the concept of T2A mutagenesis on vectors for overexpression and on a knock-in HCT116 *TP53*^+/T2A-BirA*^ cell line. In addition, we show correct localization of the BirA* enzyme in the modified cells, both before and after T2A inactivation. Based on these results, we believe that our approach is suitable for cytoplasmic and nuclear bait proteins. The current system does not allow to target the released BirA* to restricted compartments (e.g. mitochondria), which may be solved by adding localization signals. Proper localization should be carefully assessed for every bait protein, which is required for any interactome study. Our method to generate isogenic control cell lines can be readily transferred to any other proximity labeling technology (e.g. APEX ^68^). For certain bait proteins, APEX may be beneficial to characterize the temporal character of short-lived interactions, as the relative long labeling time of 16 to 24 hours in BioID does not permit for these kind of studies. However, APEX relies on exposure of the cells to hydrogen peroxide, which can be an additional source of artefacts. Therefore, it might be worthwhile to explore the use of new BioID variants such as TurboID for future proxisome analysis studies ^69^. While TurboID labeling times still exceed those of APEX (10 min compared to 1 min respectively), it is a drastic improvement over the original BirA* enzyme.

As site-directed mutagenesis is accomplished by introduction of a CRISPR/nCas9-cytidine deaminase fusion (BE3), a simple short selection step suffices to obtain a strong enrichment of cells transiently expressing the BE3 base editor and therefore having a high chance of expressing p53-MUTT2A-BirA*. In this way, there is no need to generate separate control cell lines, or to have a design wherein engineered single cell knock in clones are compared to parental populations. Such a design would inevitably lead to higher background levels as presented in this work (Figure S7, table S8). Until genome engineering progresses to efficiencies wherein clonal screening is no longer needed, T2A mutagenesis may be a valuable approach to obtain appropriate isogenic control cell lines. While the devised T2A system allows rigorous control of expression levels, also when gene expression levels are modulated by various stimuli (i.e. doxorubicin effects on p53 levels), the free BirA* cannot replicate the exact localization of the bait protein. A similar localization would allow discrimination between proteins in close proximity and interaction partners (both direct or indirect) ^70^. As with all BioID designs, downstream validation is still required to assess this aspect. While we opted for a classic well-controlled knock-in approach, novel CRISPR/Cas9 strategies would also allow the selection of isogenic populations (e.g. CRISPR co-selection ^71^, CRISPaint ^72^).

## CONCLUSIONS

Taken together, our elegant base editing approach to derive isogenic experimental cell lines from a control cell line enables BioID experiments at the endogenous level. Bringing together the advantages of proximal biotinylation in intact cells with endogenous bait expression levels and with quantitative proteomics pushes protein complex analysis to the next level and constitutes a useful addition to study hub proteins such as p53.

## Supporting information

## SUPPORTING INFORMATION

The following supporting information is available free of charge at ACS website http://pubs.acs.org.

Table S1: Primers used for cloning or screening. (PDF)

Table S2: gRNA gblock sequences cloned in pCR-Blunt vector for transfection. (PDF)

Table S3: CRISPRESSO allele frequency output. (Excel)

Table S4: GO analysis of the results from an analysis of a transient transfection p53 BioID experiment using a mock vector as control. (PDF)

Table S5: GO analysis of the results from an analysis of a transient transfection p53 BioID experiment using a bi-cistronic p53-T2A-BirA* expression construct as control. (PDF)

Table S6: Perseus data matrices generated for all analyses. (Excel)

Table S7: GO analysis of significantly enriched proteins from converted p53-MUTT2A-BirA* HCT116 cells when compared to isogenic non-converted p53-T2A-BirA* cells. (PDF)

Table S8: GO analysis of significantly enriched proteins from converted p53-MUTT2A-BirA* HCT116 cells when compared to parental HCT116 cells. (PDF)

Figure S1: Targeting strategy and primer scheme used for PCR screening. (PDF)

Figure S2: PCR screening of neomycin resistant clonal populations. (PDF)

Figure S3: PCR for assessing the removal of the selection cassette upon TAT-Cre treatment. (PDF)

Figure S4: Validation of targeted T2A-BirA* insertion. (PDF)

Figure S5: CRISPRESSO analysis of the T2A region after BE3 base editing. (PDF)

Figure S6: STRING analysis of proximal proteins for endogenous p53. (PDF)

Figure S7: Evaluation of BE3 transfected non-engineered HCT116 WT cells as control for p53 BioID. (PDF)

## ACKNOWLEDGEMENTS

We thank the VIB Proteomics Core for running the MS samples. We thank Elise Wyseure and Noortje Samyn for technical assistance. pCMV-BE3 was provided by David Liu (Addgene plasmid #73021).

## AUTHOR CONTRIBUTIONS

G.V. designed and performed cell line experiments with help of D.D.S, performed BioID experiments, analyzed data and wrote the manuscript. G.V. and D.D.S performed cell line engineering and related characterization experiments with help of A.M. J.P. operated the MS and analyzed the samples. S.E. conceived and managed the project and wrote the manuscript. K.G. is co-promotor of G.V. and read and corrected the manuscript.

