## Supplementary material for "A well-controlled BioID design for endogenous bait proteins"

#### Supplementary Information

Giel Vandemoortele[1,2], Delphine De Sutter[1,2], Aline Moliere[1,2], Jarne Pauwels[1,2], Kris Gevaert[1,2] and Sven Eyckerman[1,2,*]

[1]VIB Center for Medical Biotechnology, VIB, B-9000 Ghent, Belgium

[2]Department of Biochemistry, Ghent University, B-9000 Ghent, Belgium

*Correspondence to Prof. Dr. Sven Eyckerman

VIB Center for Medical Biotechnology

A. Baertsoenkaai 3, B-9000 Ghent, Belgium

### TABLE OF CONTENTS

Table S1: Primers used for cloning or screening. S-3
Table S2: gRNA gblock sequences cloned in pCR-Blunt vector for transfection. S-4
Table S3: Table S3. CRISPRESSO allele frequency output (column legend). S-4
Table S4: GO analysis of the results from an analysis of a transient transfection p53 BioID experiment using a mock vector as control. S-5
Table S5: GO analysis of the results from an analysis of a transient transfection p53 BioID experiment using a bi-cistronic p53-T2A-BirA* expression construct as control. S-6
Table S6: Table S6: Perseus data matrices generated for all analyses (column legend). S-7
Table S7: GO analysis of significantly enriched proteins from converted p53-MUTT2A-BirA* HCT116 cells when compared to isogenic non-converted p53-T2A-BirA* cells. S-8
Table S8: GO analysis of significantly enriched proteins from converted p53-MUTT2A-BirA* HCT116 cells when compared to parental HCT116 cells. S-9

Figure S1: Targeting strategy and primer scheme used for PCR screening. S-10
Figure S2: PCR screening of neomycin resistant clonal populations. S-12
Figure S3: PCR for assessing the removal of the selection cassette upon TAT-Cre treatment. S-13
Figure S4: Validation of targeted T2A-BirA* insertion. S-14
Figure S5: CRISPRESSO analysis of the T2A region after BE3 base editing. S-16
Figure S6: STRING analysis of proximal proteins for endogenous p53. S-17
Figure S7: Evaluation of BE3 transfected non-engineered HCT116 WT cells as control for p53 BioID. S-18

| PCR | Construct/Purpose | Forward primer (5’-3’) | Reverse primer (5’-3’) |
|---|---|---|---|
| TP53 5'HR | pAav-p53-T2A-BirA* | CACTAGGGGTTCCTGCGGCCGCAAACGCG TCAGCCCTGCACAGACATTTT | CGACGAAGCTTTGGGCCAGGATTCTCCTCG ACGTCACCGCATGTTAGCAGACTTCCTCTG CCCTCTCCACTGCCCATATGGTCTGAGTCA GGCCCTTCTG |
| TP53 3'HR | pAav-p53-T2A-BirA* | GAAGTTATGGTACCCATTCTCCACTTCTTGT TCC | GGGTTCCTGCGGCCGCTTTTGCTAGCAGAT CACGCCACTCCACTCC |
| BirA* | pAav-p53-T2A-BirA* | CATCCACGCGTCATATGGGCAGTGGAGAG GGCAGAGGAAGTCTGCTAACCTGCGGGGA CGTCGAGGAGAATCCTGGCCCAAAGGACA ACACGGTGCCCCTGAAGCTGATCGC | GCTAGCAAGCTTTCATTACTTCTCTGCGCTT CTCAGGG |
| Lox-Neo-Lox | pAav-p53-T2A-BirA* | CAGAGAAGTAATGAAAGCTTGCTAGCATA ACTTCGTATAGCATACATTATACGAAGTTA TC | GGAACAAGAAGTGGAGAATGGGTACCATA ACTTCGTATAATGTATGCTATACGAAGTTA TAGCTGGTTCT |
| TP53 junction 5'HR | Screening | ATCAGCCAAGATTGCACCAT | TCAGCTCTCCGATTCTGTCC |
| TP53 junction 3'HR | Screening | GGGAGGATTGGGAAGACAAT | CCAGTCTCCAGCCTTTGTTC |
| Cassette deletion | Cre-Lox recombination | ACTCATGTTCAAGACAGAAGGGCCTGAC | TGACGCACACCTATTGCAAGCAAGG |
| WT allele | Sanger sequencing | ACATATTTGCATGGGGTGTG | CCTAGAATGTGGCTGATTGTAAAC |
| Knock-in allele | Sanger sequencing | CACTAGGGGTTCCTGCGGCCGCAAACGCG TCAGCCCTGCACAGACATTTT | ATTACTTCTCTGCGCTTCTCAG |
| T2A region | Illumina sequencing | TCGTCGGCAGCGTCAGATGTGTATAAGAG ACAGGGGTCAGTCTACCTCCCGCC | GTCTCGTGGGCTCGGAGATGTGTATAAGA GACAGCTCGCCGTTGGCCAGCAGGG |
| Neomycin | SB probe | TGCTCCTGCCGAGAAAGTAT | GCGATGCAATTTCCTCATTT |
| T2A | pMet7 mutagenesis | CCTTTGGGCCAAAATTCTCCTCGACGTCCC CGCA | TGCGGGGACGTCGAGGAGAATTTTGGCCC AAAGG |

**Table S1. Primers used for cloning or screening**. HR: homology region; SB: Southern blot. The T2A sequence was encoded in the reverse p53 5’HR and forward BirA* primers.

| | Sequence |
|---|---|
| **gRNA1 cassette** | TGTACAAAAAAGCAGGCTTTAAAGGAACCAATTCAGTCGACTGGATCCGGTACCAAGGTCGGGCAGGAA*GAGGGCCTATTTCCCATGATTCCTTCATATTTGCATATACGATACAAGGCTGTTAGAGAGATAATTAGAATTAATTTGACTGTAAACACAAAGATATTAGTACAAAATACGTGACGTAGAAAGTAATAATTTCTTGGGTAGTTTGCAGTTTTAAAATTATGTTTTAAAATGGACTATCATATGCTTACCGTAACTTGAAAGTATTTCGATTTCTTGGCTTTATATATCTTGTGGAAAGGACGAAACACCG***ATTCTCCTCGACGTCCCCGC**GTTTTAGAGCTAGAAATAGCAAGTTAAAATAAGGCTAGTCCGTTATCAACTTGAAAAAGTGGCACCGAGTCGGTGCTTTTTTTCTAGACCCAGCTTTCTTGTACAAAGTTGGCATTA |
| **gRNA2 cassette** | TGTACAAAAAAGCAGGCTTTAAAGGAACCAATTCAGTCGACTGGATCCGGTACCAAGGTCGGGCAGGAA*GAGGGCCTATTTCCCATGATTCCTTCATATTTGCATATACGATACAAGGCTGTTAGAGAGATAATTAGAATTAATTTGACTGTAAACACAAAGATATTAGTACAAAATACGTGACGTAGAAAGTAATAATTTCTTGGGTAGTTTGCAGTTTTAAAATTATGTTTTAAAATGGACTATCATATGCTTACCGTAACTTGAAAGTATTTCGATTTCTTGGCTTTATATATCTTGTGGAAAGGACGAAACACCG***TCCTGGCCCAAAGGACAACA**GTTTTAGAGCTAGAAATAGCAAGTTAAAATAAGGCTAGTCCGTTATCAACTTGAAAAAGTGGCACCGAGTCGGTGCTTTTTTTCTAGACCCAGCTTTCTTGTACAAAGTTGGCATTA |

**Table S2. gRNA gBlock sequences cloned in pCR-Blunt® vector for transfection.** Italics: U6 promoter sequence. Bold: gRNA sequence. Underlined: universal gRNA scaffold. Sequences were ordered as IDT gBlocks and blunt-end ligated in pCR-Blunt® vectors prior to transfection in target cells together with BE3. After parallel testing it was decided to use gRNA2 for endogenous mutagenesis.

[Excel Table S3]

**Table S3. CRISPRESSO allele frequency output (column legend).** Aligned sequence: sequencing read; Reference sequence: template sequence; NHEJ: indication of Non-homologous end-joining event (NHEJ); UNMODIFIED: indication of perfect match with reference sequence (no engineering event); HDR: Homology-directed repair event (if applicable); n deleted and n inserted: nucleotides deleted or inserted compared to reference sequence; n mutated: number of nucleotides subsituted compared to reference sequence; #Reads and %Reads: absolute and relative number of reads for each sequence.

GO Cellular components

| Term | Count | P-value | Fold Enrichment | Bonferroni |
|---|---|---|---|---|
| Nucleoplasm | 380 | 8.32E-100 | 2.96 | 5.37E-97 |
| Cell-cell adherens junction | 112 | 4.24E-67 | 7.51 | 2.74E-64 |
| Cytosol | 343 | 2.48E-55 | 2.24 | 1.60E-52 |
| Cytoplasm | 422 | 2.27E-41 | 1.75 | 1.47E-38 |
| Nucleus | 417 | 2.88E-35 | 1.67 | 1.85E-32 |
| Catalytic step 2 spliceosome | 44 | 4.54E-33 | 10.36 | 2.93E-30 |
| Membrane | 220 | 2.53E-30 | 2.17 | 1.63E-27 |
| Nucleolus | 117 | 1.83E-26 | 2.96 | 1.18E-23 |
| Spliceosomal complex | 38 | 2.46E-25 | 8.76 | 1.59E-22 |
| Proteasome complex | 28 | 1.08E-20 | 10.11 | 6.94E-18 |
| Focal adhesion | 64 | 1.77E-18 | 3.55 | 1.14E-15 |
| Nuclear speck | 45 | 2.32E-18 | 4.85 | 1.49E-15 |
| Microtubule | 50 | 3.73E-14 | 3.48 | 2.41E-11 |
| Proteasome accessory complex | 13 | 4.26E-13 | 16.57 | 2.75E-10 |
| Nuclear chromosome, telomeric region | 28 | 3.57E-11 | 4.67 | 2.30E-08 |
| Eukaryotic 43S preinitiation complex | 11 | 1.00E-10 | 15.89 | 6.45E-08 |
| MLL1 complex | 14 | 1.32E-10 | 10.46 | 8.53E-08 |
| U2 snRNP | 12 | 2.15E-10 | 13.00 | 1.38E-07 |
| Actin cytoskeleton | 35 | 4.18E-10 | 3.48 | 2.70E-07 |
| Proteasome core complex | 12 | 4.32E-10 | 12.38 | 2.79E-07 |

GO Molecular function

| Term | Count | P-value | Fold Enrichment | Bonferroni |
|---|---|---|---|---|
| Poly(A) RNA binding | 256 | 5.52E-106 | 4.61 | 3.90E-103 |
| Protein binding | 679 | 4.44E-75 | 1.57 | 3.13E-72 |
| Cadherin binding involved in cell-cell adhesion | 111 | 5.31E-69 | 7.78 | 3.75E-66 |
| RNA binding | 97 | 1.30E-28 | 3.61 | 9.16E-26 |
| Unfolded protein binding | 31 | 6.35E-15 | 5.73 | 4.47E-12 |
| Transcription coactivator activity | 46 | 2.00E-14 | 3.77 | 1.41E-11 |
| Translation initiation factor activity | 21 | 4.91E-12 | 7.00 | 3.46E-09 |
| Nucleotide binding | 51 | 7.61E-12 | 2.98 | 5.37E-09 |
| Threonine-type endopeptidase activity | 12 | 8.42E-10 | 11.62 | 5.95E-07 |
| ATP binding | 126 | 1.56E-09 | 1.71 | 1.10E-06 |
| Chromatin binding | 48 | 1.29E-08 | 2.50 | 9.11E-06 |
| Histone acetyltransferase activity | 14 | 3.43E-07 | 5.93 | 2.42E-04 |
| Actin filament binding | 22 | 1.79E-06 | 3.39 | 0.001 |
| ATP-dependent RNA helicase activity | 15 | 2.50E-06 | 4.69 | 0.002 |
| RNA polymerase II distal enhancer sequence-specific DNA binding | 15 | 2.50E-06 | 4.69 | 0.002 |
| mRNA binding | 21 | 2.87E-06 | 3.42 | 0.002 |
| RNA polymerase II core binding | 9 | 3.76E-06 | 8.72 | 0.003 |
| Protein domain specific binding | 28 | 3.83E-06 | 2.74 | 0.003 |
| Ribonucleoprotein complex binding | 10 | 8.95E-06 | 6.78 | 0.006 |
| ATPase activity | 25 | 1.10E-05 | 2.78 | 0.008 |

**Table S4: GO analysis of the results from an analysis of a transient transfection p53 BioID experiment using a mock vector as control.** This table shows the top 20 enriched 'cellular component' (top panel) or 'molecular function' terms (bottom panel). Count: number of proteins identified related to a certain GO term. Bonferroni: Bonferroni-corrected p-values.

GO Cellular components

| Term | Count | P-value | Fold Enrichment | Bonferroni |
|---|---|---|---|---|
| Cytosol | 244 | 2.65E-54 | 2.64 | 1.24E-51 |
| Extracellular exosome | 211 | 2.29E-46 | 2.69 | 1.07E-43 |
| Cell-cell adherens junction | 60 | 2.20E-31 | 6.65 | 1.03E-28 |
| Focal adhesion | 63 | 2.37E-29 | 5.77 | 1.11E-26 |
| Cytoplasm | 264 | 6.74E-29 | 1.81 | 3.16E-26 |
| Myelin sheath | 40 | 8.02E-27 | 9.42 | 3.76E-24 |
| Membrane | 148 | 2.10E-25 | 2.41 | 9.82E-23 |
| Nucleoplasm | 160 | 1.60E-20 | 2.06 | 7.49E-18 |
| Extracellular matrix | 42 | 1.54E-17 | 5.08 | 7.21E-15 |
| Proteasome complex | 20 | 1.49E-15 | 11.93 | 6.75E-13 |
| Mitochondrion | 89 | 1.80E-14 | 2.39 | 8.42E-12 |
| Nucleus | 229 | 1.72E-13 | 1.51 | 8.05E-11 |
| Melanosome | 22 | 4.84E-13 | 7.80 | 2.27E-10 |
| Proteasome core complex | 12 | 1.94E-12 | 20.46 | 9.10E-10 |
| Microtubule | 34 | 4.92E-11 | 3.91 | 2.30E-08 |
| Cytosolic small ribosomal subunit | 14 | 4.17E-10 | 10.44 | 1.95E-07 |
| Mitochondrial matrix | 32 | 3.06E-09 | 3.50 | 1.43E-06 |
| Endoplasmic reticulum chaperone complex | 8 | 3.76E-09 | 26.04 | 1.76E-06 |
| Ribosome | 22 | 7.67E-09 | 4.75 | 3.59E-06 |
| Chaperonin-containing T-complex | 7 | 3.56E-08 | 27.85 | 1.67E-05 |

GO Molecular function

| Term | Count | P-value | Fold Enrichment | Bonferroni |
|---|---|---|---|---|
| Protein binding | 399 | 1.95E-39 | 1.54 | 1.27E-36 |
| Poly(A) RNA binding | 116 | 2.94E-33 | 3.48 | 1.91E-30 |
| Cadherin binding involved in cell-cell adhesion | 58 | 7.14E-31 | 6.77 | 4.65E-28 |
| Unfolded protein binding | 39 | 1.18E-30 | 11.99 | 7.67E-28 |
| ATP binding | 100 | 8.67E-15 | 2.26 | 5.65E-12 |
| Threonine-type endopeptidase activity | 12 | 3.57E-12 | 19.33 | 2.33E-09 |
| Chaperone binding | 19 | 1.73E-11 | 7.94 | 1.13E-08 |
| Structural constituent of ribosome | 30 | 1.78E-11 | 4.57 | 1.16E-08 |
| Ubiquitin protein ligase binding | 30 | 8.07E-09 | 3.54 | 5.26E-06 |
| Hsp70 protein binding | 11 | 2.29E-08 | 11.28 | 1.50E-05 |
| GTPase activity | 26 | 2.92E-08 | 3.76 | 1.90E-05 |
| GTP binding | 33 | 1.37E-07 | 2.91 | 8.91E-05 |
| Structural constituent of cytoskeleton | 17 | 1.41E-07 | 5.23 | 9.17E-05 |
| Heat shock protein binding | 11 | 2.87E-07 | 8.86 | 1.87E-04 |
| Protein kinase binding | 30 | 2.57E-06 | 2.70 | 0.002 |
| Protein complex binding | 21 | 3.19E-06 | 3.45 | 0.002 |
| Biotin carboxylase activity | 5 | 3.66E-06 | 33.83 | 0.002 |
| Adenyl-nucleotide exchange factor activity | 6 | 4.88E-06 | 20.30 | 0.003 |
| Actin filament binding | 16 | 8.20E-06 | 4.10 | 0.005 |
| Transcription coactivator activity | 22 | 1.55E-05 | 3.00 | 0.010 |

**Table S5. GO analysis of the results from an analysis of a transient transfection p53 BioID experiment using a bi-cistronic p53-T2A-BirA* expression construct as control.** This table shows the top 20 enriched 'cellular component' (top panel) or 'molecular function' terms (bottom panel). Count: number of proteins identified related to a certain cellular component. Bonferroni: Bonferroni-corrected p-values.

[Excel table S6]

**Table S6: Perseus data matrices generated for all analyses (column legend).** Each sheet corresponds to a separate analysis. Significance: significant after t-test on FDR 0.05 and optimized s0 value; -log(p-value): significance; Difference: log difference; Protein IDs, Majority protein IDs, Protein names, Gene names: protein identifiers; Peptides, Razor + unique peptides, Unique peptides: number of identified peptides for every protein; Sequence coverage, Unique + razor sequence coverage, Unique sequence coverage: % sequence coverage for every protein; Mol. Weight [kDa]: molecular weight; Q-value: adjusted p-value; Score: test score; Intensity: Observed intensities; MS/MS count: number of identifications; LFQ intensity columns: Label-Free Quantitation (LFQ) intensities for every protein in each sample.

**GO Cellular components**

| Term | Count | P-value | Fold Enrichment | Bonferroni |
|---|---|---|---|---|
| Nucleoplasm | 156 | 9.41E-87 | 5.08 | 2.25E-84 |
| Nucleus | 143 | 9.62E-34 | 2.39 | 2.30E-31 |
| Mediator complex | 21 | 6.80E-31 | 54.40 | 1.62E-28 |
| Nuclear chromatin | 26 | 7.44E-20 | 12.21 | 1.78E-17 |
| Transcription factor TFTC complex | 12 | 7.44E-20 | 77.71 | 1.78E-17 |
| STAGA complex | 11 | 1.94E-17 | 71.24 | 4.64E-15 |
| Nucleolus | 44 | 2.38E-17 | 4.65 | 5.68E-15 |
| Transcription factor TFIID complex | 14 | 4.12E-17 | 35.26 | 9.84E-15 |
| SWI/SNF complex | 9 | 1.10E-12 | 54.40 | 2.63E-10 |
| NuA4 histone acetyltransferase complex | 9 | 7.28E-12 | 45.33 | 1.74E-09 |
| NuRD complex | 9 | 7.28E-12 | 45.33 | 1.74E-09 |
| Core mediator complex | 6 | 9.01E-10 | 90.67 | 2.15E-07 |
| npBAF complex | 7 | 1.42E-09 | 52.89 | 3.39E-07 |
| nBAF complex | 7 | 4.52E-09 | 45.33 | 1.08E-06 |
| MLL1 complex | 8 | 2.20E-08 | 25.01 | 5.25E-06 |
| Protein complex | 21 | 2.97E-08 | 4.62 | 7.09E-06 |
| Sin3 complex | 6 | 1.81E-07 | 41.85 | 4.34E-05 |
| Transcription factor TFIIIC complex | 5 | 2.08E-07 | 75.56 | 4.96E-05 |
| Chromatin | 10 | 5.75E-07 | 10.19 | 1.37E-04 |
| SAGA complex | 6 | 6.00E-07 | 34.00 | 1.43E-04 |

**GO Molecular function**

| Term | Count | P-value | Fold Enrichment | Bonferroni |
|---|---|---|---|---|
| Transcription coactivator activity | 40 | 1.48E-32 | 13.41 | 3.88E-30 |
| Protein binding | 179 | 3.12E-28 | 1.69 | 8.19E-26 |
| RNA polymerase II transcription cofactor activity | 18 | 7.44E-24 | 41.58 | 1.96E-21 |
| Histone acetyltransferase activity | 19 | 6.15E-23 | 32.92 | 1.62E-20 |
| Chromatin binding | 34 | 6.31E-19 | 7.23 | 1.66E-16 |
| Nucleosomal DNA binding | 15 | 1.31E-16 | 27.12 | 2.92E-14 |
| RNA polymerase II distal enhancer sequence-specific DNA binding | 16 | 1.04E-15 | 20.47 | 2.63E-13 |
| DNA binding | 61 | 2.41E-15 | 3.03 | 6.42E-13 |
| Histone deacetylase activity | 12 | 6.86E-13 | 25.59 | 1.80E-10 |
| Vitamin D receptor binding | 9 | 2.19E-12 | 49.89 | 5.76E-10 |
| Poly(A) RNA binding | 43 | 3.18E-11 | 3.17 | 8.37E-09 |
| p53 binding | 12 | 3.93E-10 | 14.89 | 1.03E-07 |
| Thyroid hormone receptor binding | 9 | 6.68E-10 | 27.72 | 1.76E-07 |
| Transcription cofactor activity | 12 | 7.50E-10 | 14.05 | 1.97E-07 |
| RNA polymerase II core promoter proximal region sequence-specific DNA binding | 20 | 5.53E-08 | 4.68 | 1.45E-05 |
| Transcription factor binding | 18 | 5.76E-08 | 5.27 | 1.51E-05 |
| RNA polymerase II repressing transcription factor binding | 7 | 6.54E-07 | 21.56 | 1.72E-04 |
| Lysine-acetylated histone binding | 6 | 1.76E-06 | 27.72 | 4.63E-04 |
| Ligand-dependent nuclear receptor transcription coactivator activity | 8 | 2.35E-06 | 13.04 | 6.17E-04 |
| Ligand-dependent nuclear receptor binding | 6 | 4.07E-06 | 23.76 | 0.001 |

**Table S7: GO analysis of significantly enriched proteins from converted p53-MUTT2A-BirA* HCT116 cells when compared to isogenic non-converted p53-T2A-BirA* cells.** This table shows the top 20 enriched ‘cellular component ’(top panel) or ‘molecular function’ terms (bottom panel). Count: number of proteins identified related to a certain GO term. Bonferroni: Bonferroni-corrected p-values.

**GO Cellular components**

| Term | Count | P-value | Fold Enrichment | Bonferroni |
|---|---|---|---|---|
| Nucleoplasm | 252 | 1.48E-103 | 4.03 | 5.75E-101 |
| Nucleus | 264 | 8.52E-49 | 2.17 | 3.32E-46 |
| Nucleolus | 89 | 1.73E-34 | 4.63 | 6.75E-32 |
| Catalytic step 2 spliceosome | 33 | 7.58E-30 | 15.98 | 2.95E-27 |
| Mediator complex | 23 | 3.19E-28 | 29.28 | 1.24E-25 |
| Cytosolic large ribosomal subunit | 25 | 7.53E-23 | 16.38 | 2.93E-20 |
| Nuclear speck | 37 | 2.15E-22 | 8.20 | 8.36E-20 |
| Nuclear chromatin | 31 | 4.50E-17 | 7.16 | 1.75E-14 |
| Ribosome | 29 | 5.73E-17 | 7.78 | 4.32E-14 |
| Transcription factor TFTC complex | 12 | 2.11E-16 | 38.19 | 8.64E-14 |
| Cell-cell adherens junction | 38 | 2.66E-16 | 5.24 | 8.64E-14 |
| Spliceosomal complex | 21 | 1.87E-14 | 9.95 | 7.26E-12 |
| STAGA complex | 11 | 2.62E-14 | 35.01 | 1.02E-11 |
| Cytoplasm | 188 | 7.43E-14 | 1.60 | 2.89E-11 |
| Membrane | 103 | 2.81E-13 | 2.09 | 1.09E-10 |
| Transcription factor TFIID complex | 14 | 4.25E-13 | 17.33 | 1.65E-10 |
| NuA4 histone acetyltransferase complex | 11 | 1.06E-12 | 27.23 | 4.11E-10 |
| Focal adhesion | 35 | 1.49E-11 | 3.99 | 5.80E-09 |
| MLL1 complex | 12 | 1.49E-11 | 18.44 | 5.80E-09 |
| U2 snRNP | 10 | 1.75E-10 | 22.28 | 6.80E-08 |

**GO Molecular function**

| Term | Count | P-value | Fold Enrichment | Bonferroni |
|---|---|---|---|---|
| Poly(A) RNA binding | 148 | 6.17E-70 | 5.41 | 2.67E-67 |
| Protein binding | 346 | 1.23E-44 | 1.63 | 5.31E-42 |
| Transcription coactivator activity | 47 | 1.15E-27 | 7.82 | 4.99E-25 |
| RNA polymerase II transcription cofactor activity | 20 | 7.30E-22 | 22.93 | 3.15E-19 |
| RNA binding | 57 | 2.61E-20 | 4.30 | 1.13E-17 |
| Histone acetyltransferase activity | 19 | 1.98E-17 | 16.34 | 8.54E-15 |
| Cadherin binding involved in cell-cell adhesion | 38 | 8.49E-17 | 5.41 | 4.80E-14 |
| Vitamin D receptor binding | 12 | 1.79E-15 | 33.02 | 7.67E-13 |
| Structural constituent of ribosome | 31 | 1.89E-14 | 5.76 | 8.15E-12 |
| Chromatin binding | 40 | 5.77E-14 | 4.22 | 2.49E-11 |
| Nucleosomal DNA binding | 16 | 1.12E-13 | 14.36 | 4.85E-11 |
| DNA binding | 89 | 1.31E-12 | 2.19 | 5.64E-10 |
| RNA polymerase II distal enhancer sequence-specific DNA binding | 17 | 2.22E-12 | 10.79 | 9.61E-10 |
| Thyroid hormone receptor binding | 12 | 1.32E-11 | 18.34 | 5.72E-09 |
| Transcription cofactor activity | 15 | 1.28E-09 | 8.72 | 5.52E-07 |
| Histone deacetylase activity | 12 | 1.31E-09 | 12.70 | 5.67E-07 |
| Transcription factor binding | 26 | 2.79E-08 | 3.78 | 1.20E-05 |
| p53 binding | 13 | 6.11E-08 | 8.01 | 2.64E-05 |
| Nucleotide binding | 27 | 3.86E-07 | 3.20 | 1.67E-04 |
| RNA polymerase II core promoter proximal region sequence-specific DNA binding | 26 | 1.91E-06 | 3.02 | 8.25E-04 |

**Table S8: GO analysis of significantly enriched proteins from converted p53-MUTT2A-BirA* HCT116 cells when compared to parental HCT116 cells.** This table shows the top 20 enriched 'cellular component' (top panel) or 'molecular function' terms (bottom panel). Count: number of proteins identified related to a certain GO term. Bonferroni: Bonferroni-corrected p-values.

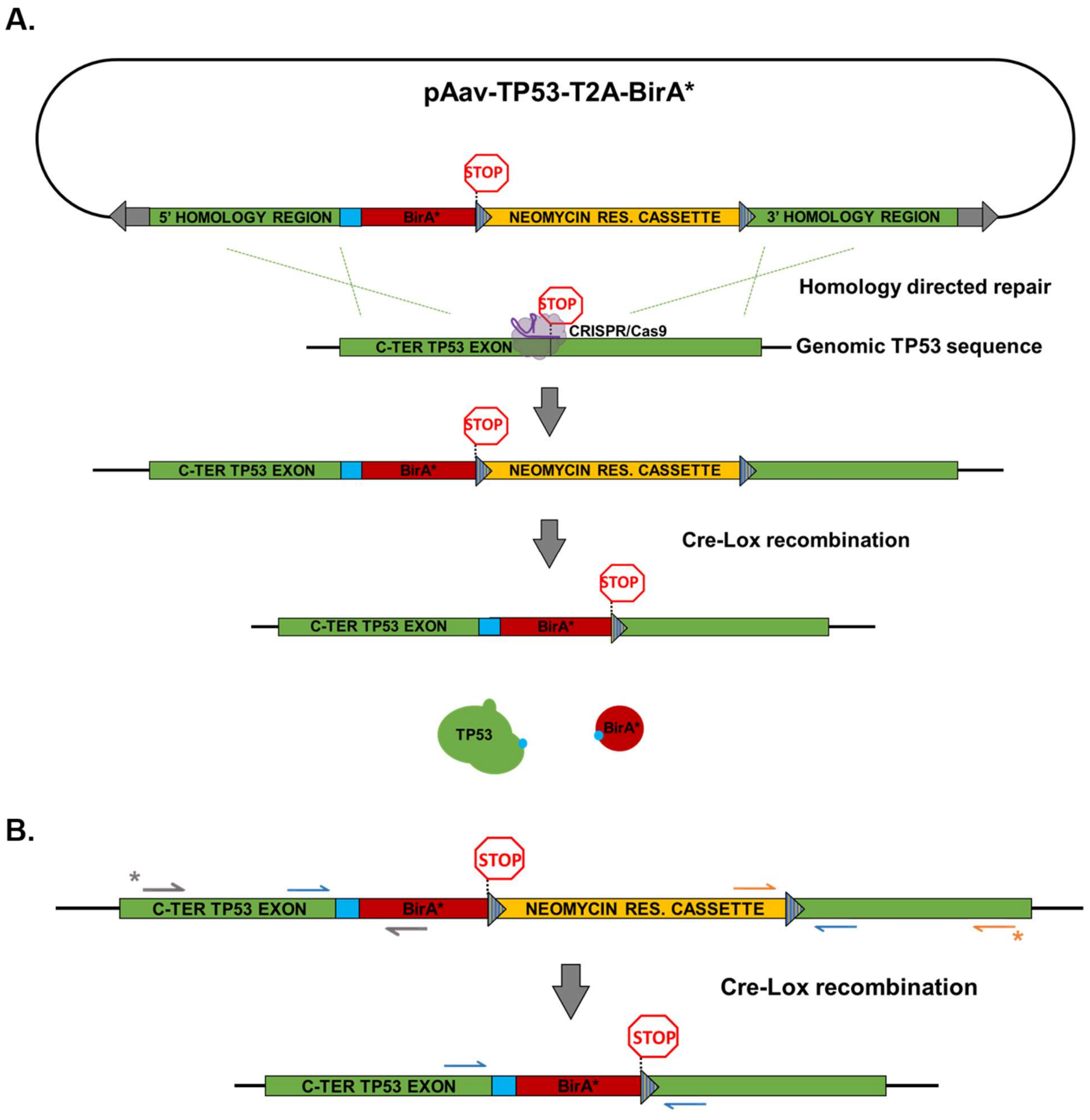


**Figure S1. Targeting strategy and primer scheme used for PCR screening.** (A) Knock-in of T2A-BirA* to the p53 C-terminus. The pAav-TP53-T2A-BirA* plasmid was packaged in rAAV and served as HDR repair template for the Cas9-induced DNA lesion site. After a first screening round to confirm integration, cells were treated with TAT-Cre for Cre-Lox recombination leading to the removal of the neomycin resistance cassette. The final HCT116 *TP53*$^{+/T2A\text{-}BirA*}$ cell line contains only a remnant Lox site as a scar sequence. Cyan:

T2A sequence. Purple: CRISPR/Cas9. (B) PCR screening strategy. Arrows are indicative for primer location on the altered locus for PCR screening. Grey arrows: primers used for 5' HR PCR (1706 nt). Orange arrows: primers used for 3' HR PCR (1467 nt). Blue arrows: primers used for cassette deletion PCR (WT amplicon: 163 nt; recombined amplicon: 1230 nt; not recombined amplicon: 2879 nt). Asterisks indicate location of the primer outside the HR to exclude false-positive PCR amplicon in case of random integration.

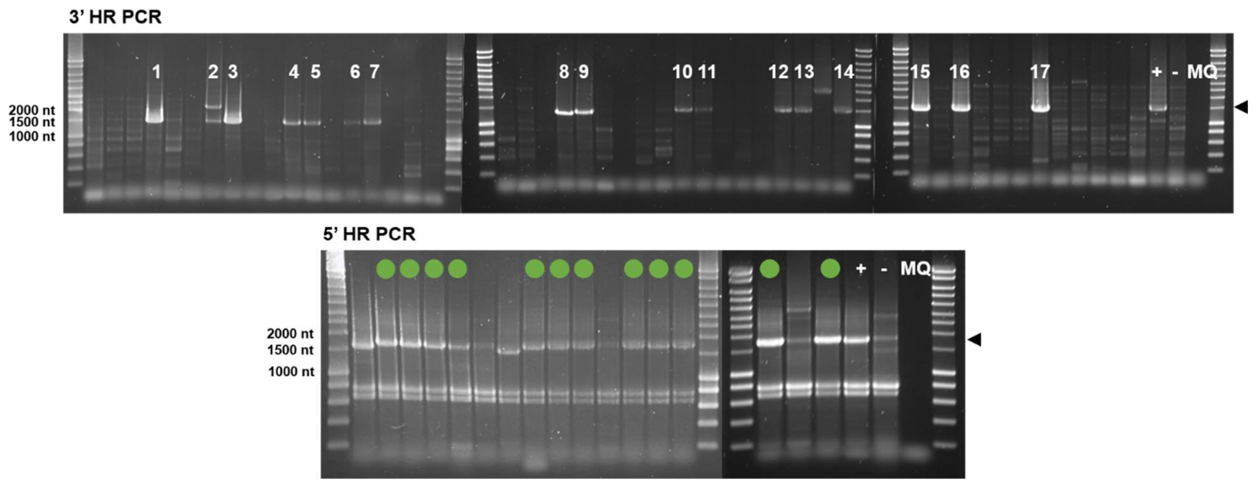


**Figure S2. PCR screening of neomycin resistant clonal populations.** PCR screening of clonal populations for HDR on agarose gels. Lysates were obtained by adding the cell suspension to DirectPCR lysis reagent. 17 clones showed a correct amplicon for 3' HR (1467 nt). These 17 clones were retained for the 5' HR PCR of which 12 were positive for the 5' HR PCR (1706 nt) as indicated in green. +: Positive control consisting of the neomycin resistant pool of HCT116 cells subjected to transfection and infection. -: Parental (non-engineered) HCT116 cells. MQ: No input DNA negative control.

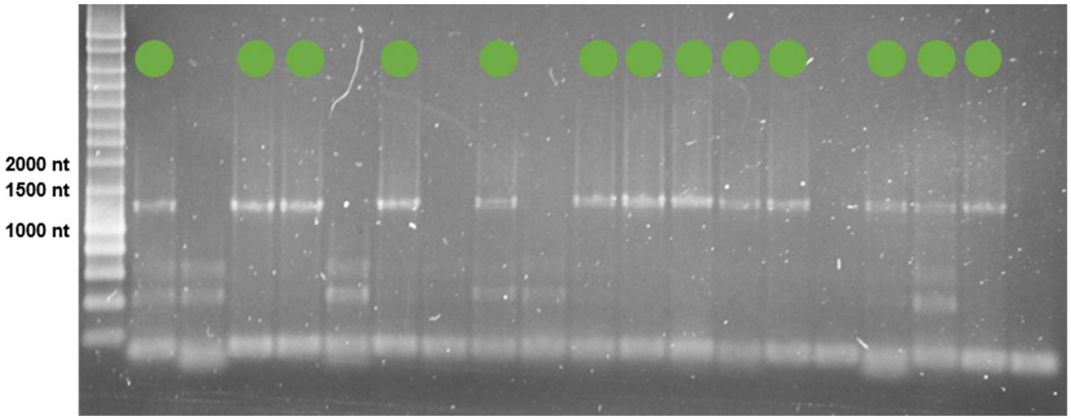


**Figure S3. PCR for assessing the removal of the selection cassette upon TAT-Cre treatment.** Agarose gel showing a typical post-recombination PCR screening. Lysates were obtained by adding the cell suspension to DirectPCR lysis reagent. Confirmed HDR positive clones were subjected to TAT-Cre treatment and screened by PCR for neomycin selection cassette removal by Cre-Lox recombination. WT amplicon: 163 nt; recombined amplicon: 1230 nt; not recombined amplicon: 2879 nt. 13 out of 19 clones showed recombination as indicated in green. Contrast adjusted to +40%. Empty lanes are possibly due to not-recombined amplicons that were not amplified as elongation time did not allow for PCR amplification of this size. Alternatively, this can be due to varying amounts of input DNA obtained from the lysis reagent.

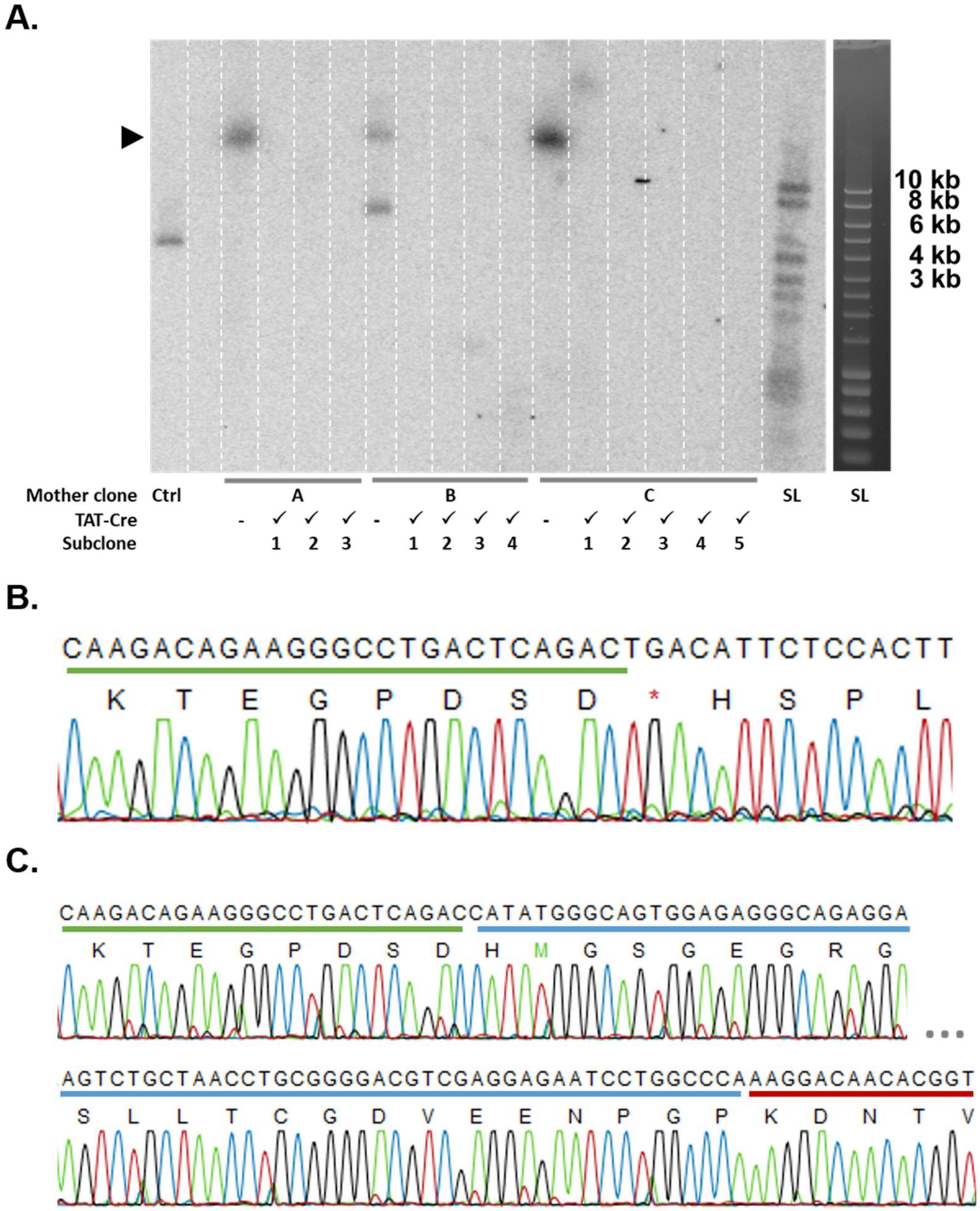


**Figure S4. Validation of targeted T2A-BirA* insertion.** (A) Southern blot of engineered T2A-BirA* clones prior and after TAT-Cre treatment to verify single integration and subsequent removal of the neomycin selection cassette. A total of 3, 4 and 5 TAT-Cre treated single cell subclones derived from 3 mother clones (A, B and C respectively) were assayed for cassette removal. A [α-$^{32}$P] Southern blot probe was generated against the neomycin resistance cassette. Expected size of the EcoRI digested HCT116 *TP53*$^{+/\text{T2A-BioID}}$ DNA fragment containing the probed selection cassette: ±20 kb. Ctrl: positive control from an ERN1 FLAG-tagged cell line containing the cassette with expected fragment size of 3537 bp. SL: Smartladder (left: post transfer on the membrane, right: SL on 0.7% agarose gel prior to transfer). Mother clone B showed double integration of the selection cassette, leading to exclusion of all derived post TAT-Cre expanded clones for further experiments. Clones 1-4 in the T2A mutagenesis Western blot experiment (Figure 2A) correspond to clone A1, A3, C1 and C3 respectively. (B) Sanger sequencing result of unaltered WT allele in knock-in clone C3 (clone 4). Green line: C-terminus of p53 (C) Sanger sequencing result of T2A-BirA* allele of the HCT116 $TP53^{+/\text{T2A-BirA*}}$ clone C3 (clone 4). Blue line: Linker sequence and T2A sequence. Red line: N-terminus of BirA*

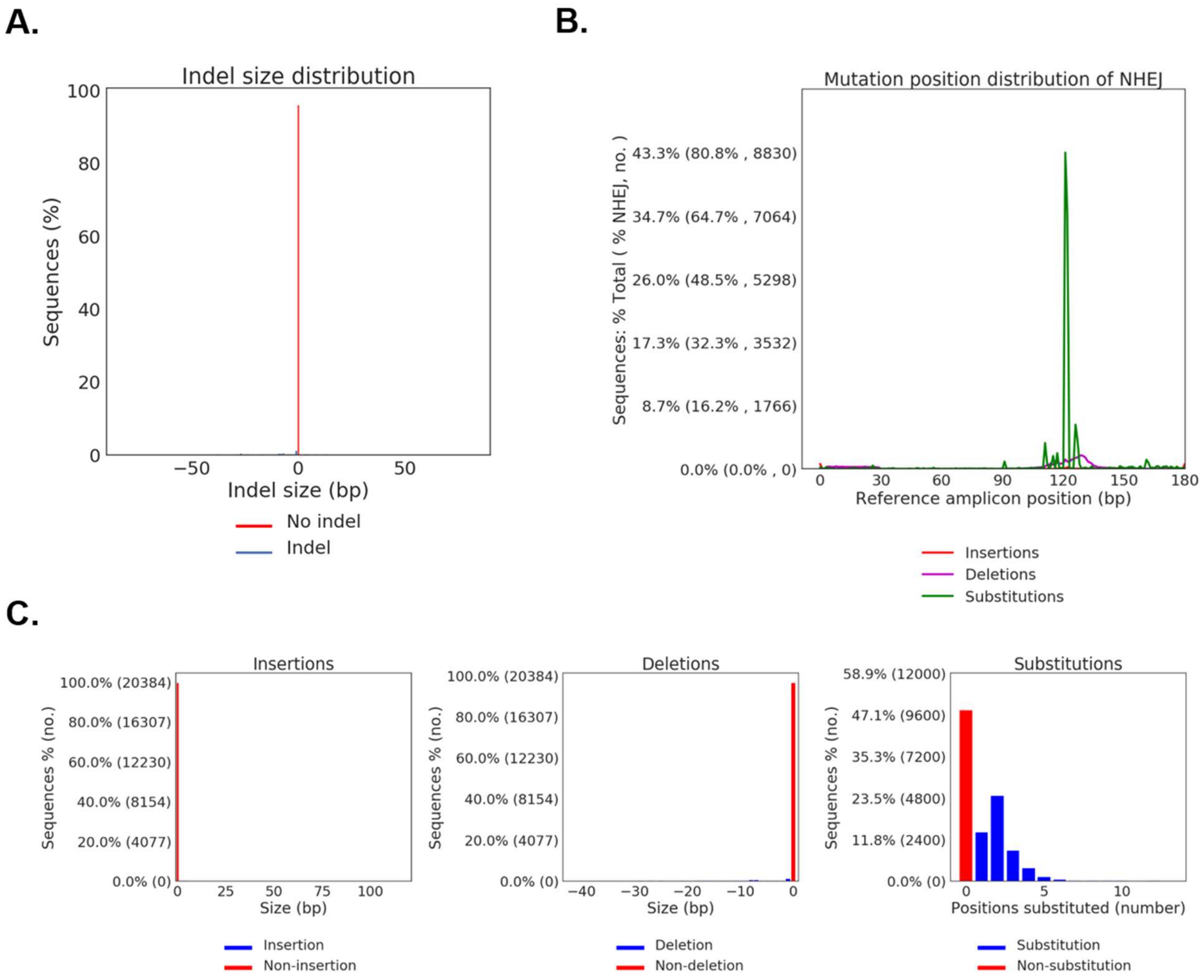


**Figure S5. CRISPRESSO analysis of the T2A region after BE3 base editing.** (A) Insertion and deletion (Indel) size distribution after BE3 treatment. Formation of indel mutations was negligible, with nearly all mutated amplicons bearing substitutions. (B) Position distribution of mutations in BE3-treated cells. Substitutions are exclusively prevalent in the mutation window of BE3. (C) Mutational size pattern of BE3-treated cells. Negligible insertions (left panel) and deletions (middle panel) were discerned. Substitutions (right panel) vary in size from 1 to 5 nucleotides, in accordance with the number of available residues in the T2A target sequence.

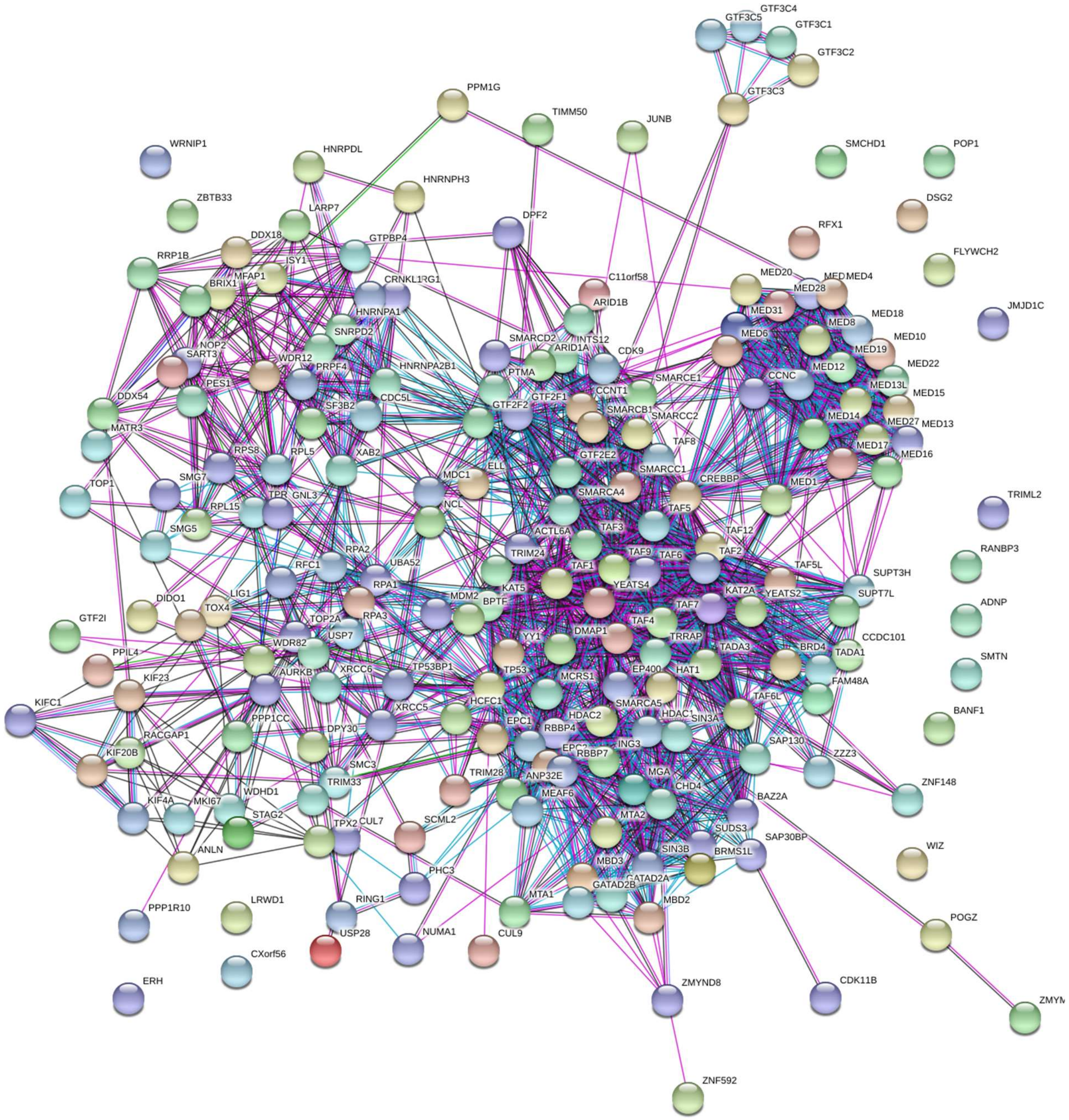


**Figure S6. STRING analysis of proximal proteins for endogenous p53.** All candidate proximal proteins detected in the edited p53-MUTT2A-BirA* HCT116 cells when isogenic p53-T2A-BirA* cells were used as control. Text mining associations are disabled. Minimum required interaction score was set on medium confidence (0.400).

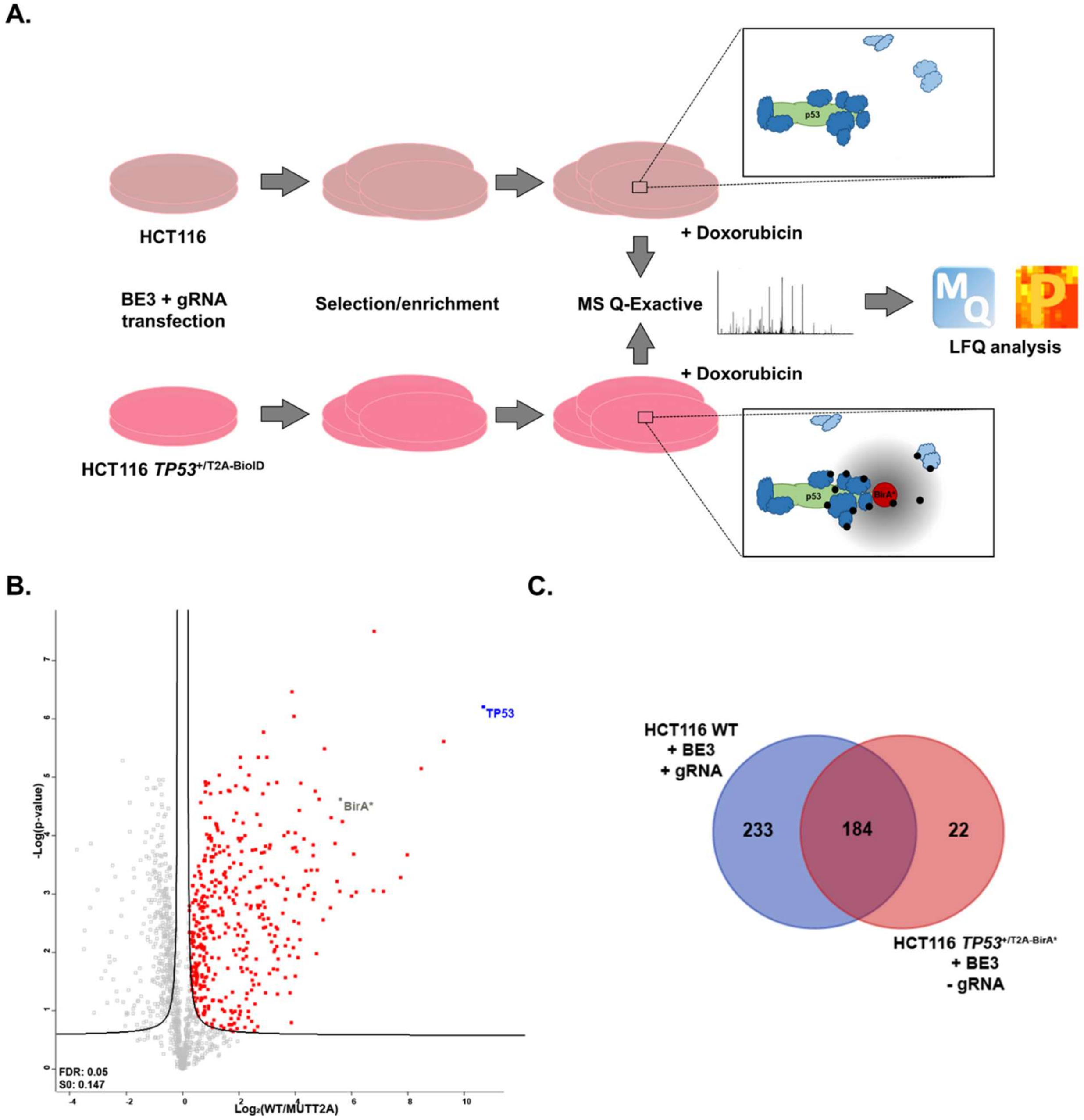


**Figure S7. Evaluation of BE3 transfected non-engineered WT cells as control for p53 BioID.** (A) Proteomics workflow when using transfected HCT116 WT cells as control condition. (B) Volcano plot showing the differential LFQ intensity levels (X-axis) and the p-values (Y-axis) for HCT116 *TP53*$^{+/\text{T2A-BirA*}}$ after T2A conversion in presence of 1μM doxorubicin. (C) Venn diagram comparing significant hits in analyses using the two different control conditions. Comparing the two different controls showed an overlap of 184 significant proteins. 233 proteins were exclusively identified in a significant manner when using the HCT116 BE3-transfected control.
